# Conditional Antibody Expression to Avoid Central B Cell Deletion in a Humanized HIV-1 Vaccine Mouse Models

**DOI:** 10.1101/2020.01.03.894279

**Authors:** Ming Tian, Kelly McGovern, Hwei-Ling Cheng, Peyton Waddicor, Lisa Rieble, Mai Dao, Yiwei Chen, Michael T. Kimble, Elizabeth Cantor, Nicole Manfredonia, Rachael Judson, Aimee Chapdelaine-Williams, Derek W. Cain, Barton F. Haynes, Frederick W. Alt

**Affiliations:** Program in Cellular and Molecular Medicine, Boston Children’s Hospital, Department of Genetics, Harvard Medical School, Howard Hughes Medical Institute, Boston, MA02115, USA; Department of Medicine, Duke Human Vaccine Institute, Duke School of Medicine, Durham, NC 27710 USA; Department of Immunology, Duke University School of Medicine, Durham, NC 27710, USA

**Keywords:** HIV-1, immunoglobulin, mouse model

## Abstract

HIV-1 vaccine development aims to elicit broadly neutralizing antibodies (bnAbs) against diverse viral strains. In some HIV-1 infected individuals, bnAbs evolve from precursor antibodies through affinity maturation. To induce bnAbs, a vaccine must mediate a similar process of antibody maturation. One way to test vaccination strategies is to immunize mouse models that express human bnAb precursors. Such immunization experiments can assess whether the vaccine can convert precursor antibody into bnAb. A major problem with such mouse models is that bnAb expression often hinders B cell development in the bone marrow. Such developmental blocks may be attributed to unusual properties of bnAb variable regions, such as poly-reactivity and long antigen-binding loops, which are often under negative selection during primary B cell development. To address this problem, we devised a method to circumvent B cell developmental block by expressing bnAbs conditionally in mature B cells. We validated this method by expressing the unmutated common ancestor (UCA) of the human VRC26 bnAb in transgenic mice. Constitutive expression of combined immunoglobulin heavy and light chains of VRC26UCA led to developmental arrest of B cell progenitors in the bone marrow; poly-reactivity of VRC26UCA and poor pairing of VRC26UCA IgH chain with mouse surrogate light chain may contribute to the phenotype. The conditional expression strategy circumvented this developmental impediment, allowing the VRC26UCA to be expressed in mature peripheral B cells. This method should be generally applicable for expressing other antibodies that are under negative selection during B cell development.

**Significance statement:** Mouse models can provide fast and cost-effective systems to test HIV-1 vaccine candidates at the pre-clinical stage. To serve this purpose, mouse models are engineered to express the precursors of human bnAbs against diverse HIV-1 strains. Immunization of such mouse models can evaluate the ability of vaccines to mature the precursor antibodies into bnAbs. However, due to unusual properties of bnAbs, mouse models expressing their precursors often have B cell developmental defects. In this study, we devised and validated a strategy to address this problem. This method could also facilitate the expression of other clinically relevant antibodies in mature B cells in transgenic mice; immunization of such mice could be used to generate novel antibodies with desirable properties.

## Introduction

Some HIV-1 infected individuals develop broadly neutralizing antibodies (bnAbs) against diverse viral strains (1, 2). A goal of HIV-1 vaccine development is to elicit comparable bnAbs. The bnAb developmental pathway during natural infection can serve as a blueprint for immunogen design. In bnAb donors, each bnAb lineage evolves from a germline antibody precursor, or unmutated common ancestor (UCA). The maturation of the UCA into a bnAb entails extensive somatic hypermutation. Conversion of UCA into bnAbs via vaccination may require a series of immunogens to guide the maturation process. These immunogens can be tested in transgenic mice that express bnAb precursor (3).

To express bnAb precursors in transgenic mice, a standard method is to integrate, or knock-in (KI), the pre-assembled variable region (V(D)J) exons of the immunoglobulin (Ig) heavy (HC) and light chain (LC) of the antibody into mouse J_H_ and Jκ loci, respectively (3). By allelic exclusion, pre-assembled V(D)J exons inhibit rearrangement of the endogenous Ig loci (4), and, in theory, all B cells in the transgenic mice should express the KI antibody. A common problem with such KI mice is that B cells expressing certain bnAbs are deleted or anergized (5–10). Such phenotypes are characteristic of B cells expressing auto-reactive antibodies. Some bnAbs are indeed poly-reactive or auto-reactive (11–16). Phenomena known as V_H_ replacement and IgL receptor editing can allow selection of B cells that have deleted the HC and/or LC V(D)J exons of autoreactive antibodies and replaced them with endogenously assembled V(D)J exons (17–22). Clonal deletion or anergy can eliminate or inactivate autoreactive B cells (23–27). Exclusion from B cell follicles precludes autoreactive B cells from the compartment of recirculating mature B cells (28, 29). These tolerance control mechanisms hinder the expression of bnAbs in KI mice (30).

We have devised a method to overcome the developmental restriction on BnAb V(D)J KI alleles. For this purpose, we expressed a human bnAb UCA conditionally in mature B cells, thereby circumventing tolerance control checkpoints during B cell maturation. As a proof-of-principle experiment, we used this method to express the UCA of human VRC26 bnAb (31, 32). The VRC26 bnAb interacts with the V1V2 region at the apex of HIV-1 Envelop protein. The HC of VRC26 features an extraordinarily long complementarity determining region 3 (CDR H3), consisting of 37 amino acids. The long CDR H3 penetrates through thick glycan layer on HIV-1 Envelop to interact with a conserved peptide epitope underneath (33). Antibodies with such a long CDR H3 is extremely rare in both human and mouse (34, 35). Antibodies with long CDR H3s are often subject to negative selection during B cell development (36). We anticipated similar obstacles for VRC26 expression in mice and confirmed this to be the case, allowing us to study the underlying cause and test our approach to address the problem.

## Results

We constructed two mouse models for this study, constitutive and conditional expression models. In the constitutive expression model, V(D)J exons of the VRC26UCA HC and LC were integrated into mouse J_H_ and Jκ locus, respectively (Fig.1A; Fig.S1). This conventional KI mouse could reveal negative selection against VRC26UCA and serve as a reference point for the conditional expression model. The design of the conditional expression model was adapted from a method to switch antibody V regions in memory B cells (37). In this model, two V(D)J exons were integrated in tandem into the mouse J_H_ locus (Fig.1B; Fig.S1). The V(D)J exon proximal to the μ constant region (Cμ) derives from the IgH chain of a mouse antibody that promotes normal B cell development (38). The intronic enhancer (iEμ), immediately downstream of the mouse V(D)J exon, should preferentially activate the expression of the mouse heavy chain (HC) over the VRC26UCA HC V exon inserted upstream. In addition, two polyadenylation sites truncate transcripts from the upstream VRC26UCA V exon (Fig.1C). These two measures ensure that the mouse HC is expressed in B cell progenitors and drives B cell maturation, without interference from VRC26UCA. For this reason, we refer to the mouse antibody as the “driver”. Two loxP sites flank the driver V exon and polyadenylation sites (Fig.1C). The mouse model also harbors a CD21-cre transgene (39), which becomes active in mature B cells and excises the driver V(D)J exon and the polyadenylation sites. Consequently, VRC26UCA replaces the driver for expression in mature B cells. This strategy should enable VRC26UCA expression to bypass the B cell maturation block. The conditional expression method is applicable for both HC and LC. In the case of VRC26UCA, we hypothesized that the long CDR H3 would be the primary target for negative selection. To test this idea, we expressed the HC of VRC26UCA conditionally, but the LC constitutively (Fig.1B). In this situation, phenotypic differences between the conditional and constitutive models can be ascribed to differential HC expression.

**Fig. 1.**
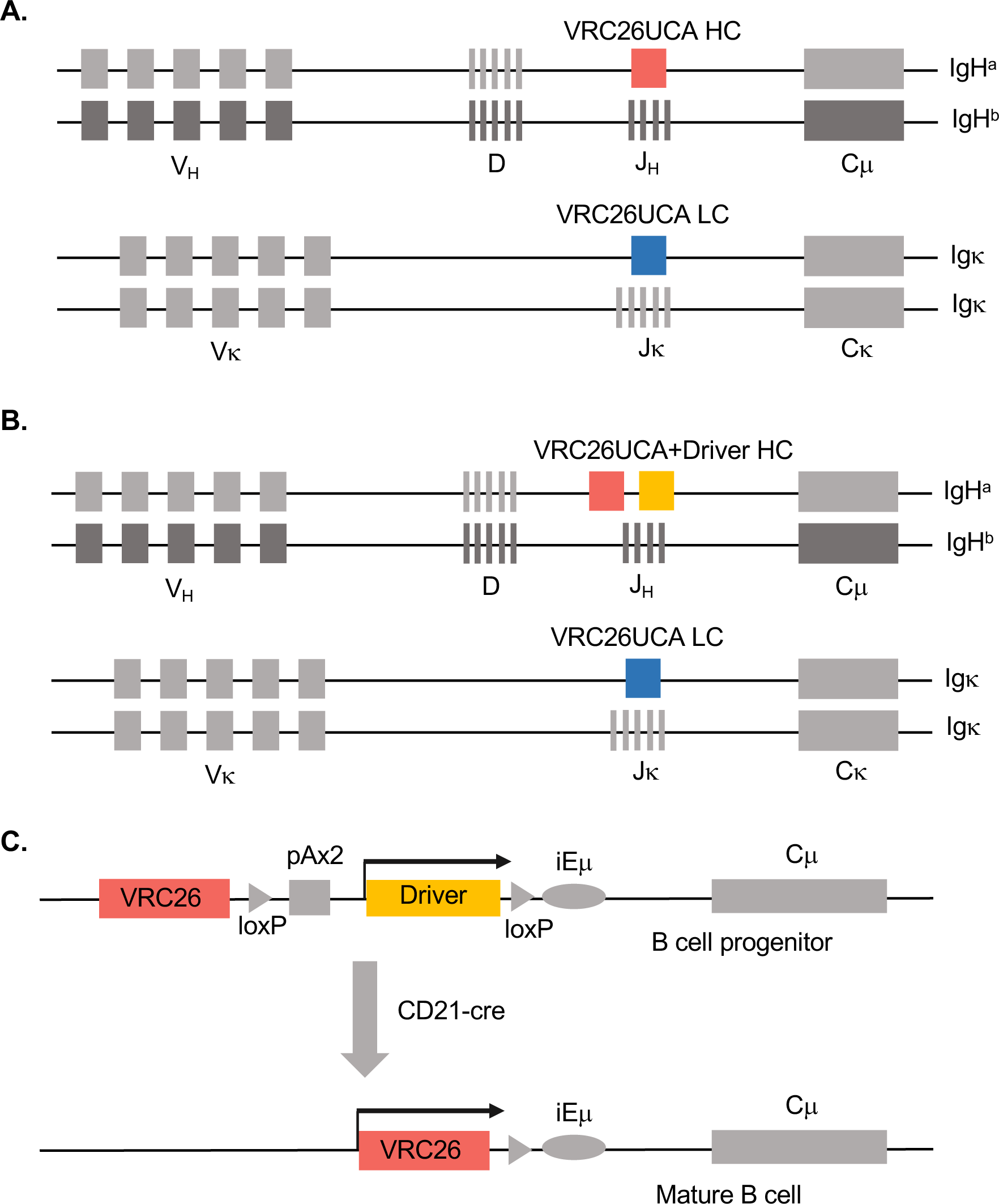
Diagram to illustrate constitutive and conditional expression model of VRC26UCA. **(A)** Illustration of constitutive expression model of VRC26UCA. The diagram shows the two IgH allele and Igκ alleles. V_H_, D, J_H_ gene segments and Cμ exons are marked on IgH; Vκ, Jκ gene segments and Cκ exon are marked on Igκ. The diagram is for illustration purpose only and is not drawn to scale. In the constitutive expression model of VRC26UCA, pre-rearranged V exons of VRC26UCA HC and LC were integrated into the mouse JH and Jκ locus, respectively. The integration was performed in a F1 ES cell line that was derived from 129Sv and C57BL/6 mouse strains. VRC26UCA HC was integrated into the IgH^a^ allele from 129Sv mouse strain. The other IgH^b^ allele from C57BL/6 mouse strain was unmodified. The two Igκ alleles are not distinguishable by allotypic markers. **(B)** Illustration of conditional expression model of VRC26UCA. In this model, a conditional expression cassette was integrated into the J_H_ locus of IgH^a^ allele. The cassette consists of two pre-rearranged V exons in tandem, which encode the V regions of VRC26UCA HC and driver HC (VRC26UCA+driver HC), respectively. A pre-rearranged VRC26UCA LC V exon was integrated into Jκ locus. **(C)** Illustration of the conditional expression cassette. Two factors contributed to the preferential expression of the driver HC in B cell progenitors. First, the V exon for the driver HC was positioned closer to the intronic enhancer (iEμ) and Cμ, than the V exon for VRC26UCA HC. Second, two polyadenylation sites (pAx2) truncated transcripts from the upstream VRC26UCA V exon. In mature B cells, CD21-cre in the mouse model became active. Recombination between the two loxP sites deleted the two polyadenylation sites and driver HC. As a result, VRC26UCA HC was expressed in mature B cells.

### B cell deletion in the constitutive expression model of VRC26UCA

We used allotypic markers to monitor VRC26UCA expression in the constitutive expression model. The two IgH alleles of the mouse model were derived from 129Sv and C57BL/6 mouse strains, respectively. The KI VRC26UCA HC resided on the IgH^a^ allele from 129Sv strain; the IgH^b^ allele from C57BL/6 strain was unmodified (Fig.1A). In naïve B cells, VRC26UCA HC was expressed as IgM^a^, whereas mouse HC was expressed as IgM^b^. In wild-type F1 mice, equal proportions of splenic B cells expressed IgM^a^ or IgM^b^ (Fig.2A). Expression of pre-rearranged V exons from IgH^a^ allele should allelically exclude the IgH^b^ allele; consequently, most B cells in such KI mice should express IgM^a^. The B1-8/3-83 KI mouse exemplified this phenomenon (Fig.2B). This mouse line harbors pre-rearranged V(D)J exons for the B1-8HC and 3-83LC at the J_H_ and Jκ loci, respectively (40, 41). B1-8 is a mouse antibody against 4-hydroxy-3-nitrophenylacetyl hapten (42, 43). 3-83 is a mouse antibody against H-2^k^ class I MHC (44). The combination of B1-8HC and 3-83LC forms an innocuous antibody that supports B cell development (45). Splenic B cells from B1-8/3-83 KI mouse expressed exclusively IgM^a^ (Fig.2B). By contrast, splenic B cells in the constitutive expression model of VRC26UCA expressed predominantly IgM^b^. In 7-week old mice, some IgM^a^ B cells were detectable, but their surface IgM^a^ levels were abnormally low (Fig.2C). By 37-weeks of age, even fewer B cells were IgM^a+^ (Fig.2D). Another sign of a B cell anomaly was that VRC26UCA HC/LC mouse had reduced numbers of B cells in the spleen relative to F1 mouse (compare Fig.S2E and S2F with S2C). Using T cell numbers as an internal reference, the B/T ratio in F1 mouse was nearly 1.1; the ratio was reduced to 0.28 in 7-week old VRC26UCA HC/LC mouse. A KI mouse with VRC26UCA HC alone had a similar, but less severe, phenotype (Fig.2I and 2J, Fig.S2G and S2H). On the other hand, a KI mouse with only the VRC26UCA LC had nearly normal B cell numbers (Fig.S2I and S2K).

**Fig. 2.**
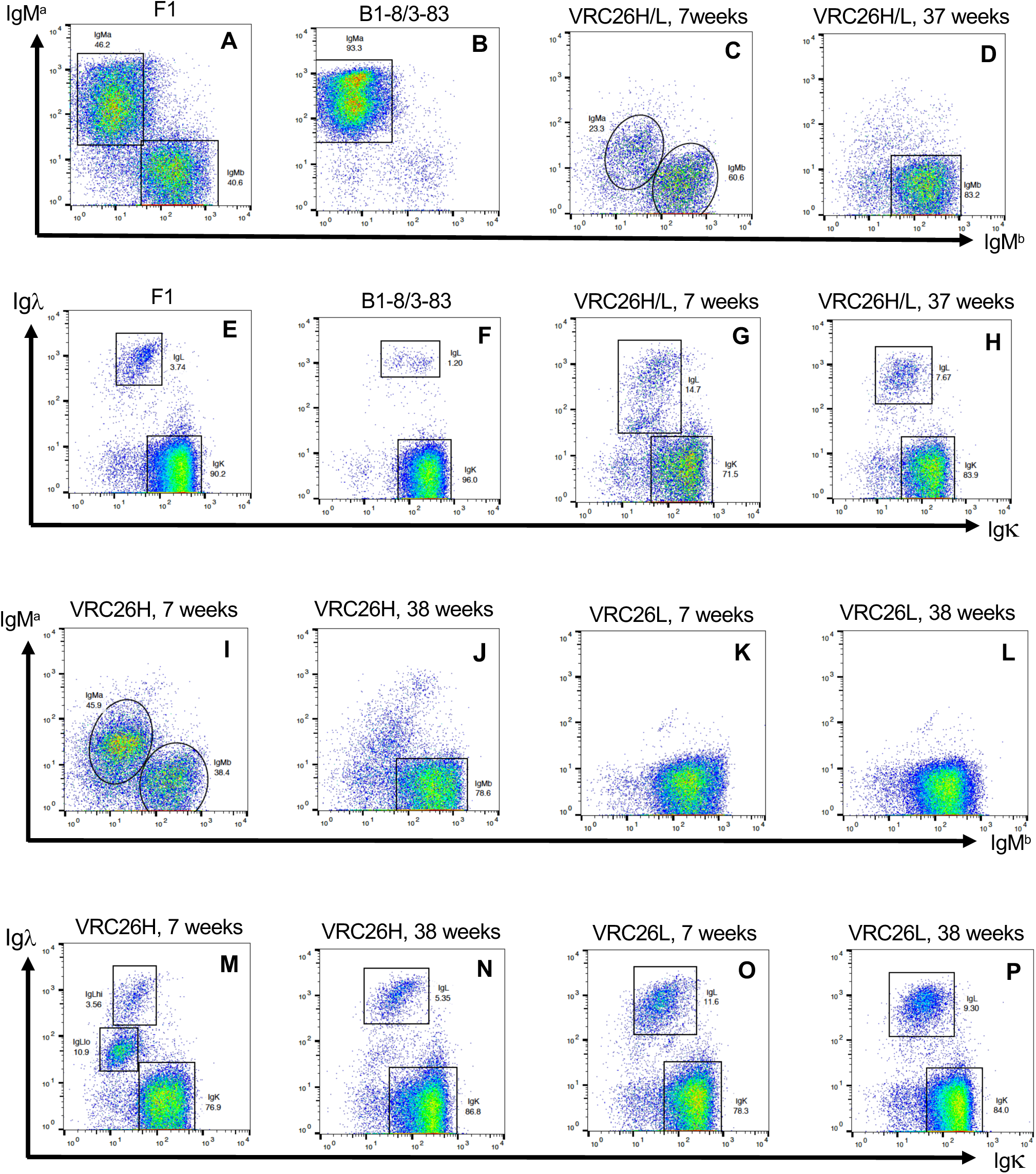
FACS analysis of splenic B cells in the constitutive expression model of VRC26UCA. All the plots were gated on B cells. Gating strategy was illustrated in Fig.S2. The staining antibodies were indicated next to the axis. **(A-D; I-L)** These plots show staining patterns of IgM^a^ versus IgM^b^. **(E-H; M-P)** These plots show staining patterns of Igκ versus Igλ. Mouse genotype was shown above each plot: F1, 129Sv x C57BL/6 wild-type mouse; B1-8/3-83, KI mice with B1-8HC and 3-83LC; VRC26H/L, constitutive expression model of VRC26UCA HC and LC; VRC26H, KI mice for constitutive expression of VRC26UCA HC; VRC26L, KI mice for constitutive expression of VRC26UCA LC. The ages of the VRC26UCA KI mice were indicated above the plot.

The pattern of Ig light chain expression in the VRC26UCA KI mouse was also unusual. Unlike IgH, Igκ alleles in this model are not distinguishable by allotypic markers. Igκ^+^ B cells could express either VRC26UCA LC or a mouse LC, but all Igλ^+^ B cells expressed mouse LC. In normal mice, about 3-5% of B cells expressed Igλ, as shown in F1 mouse (Fig.2E). Because of allelic exclusion, expression of pre-rearranged VJ exon from the κ locus, as in B1-8/3-83 KI mouse, further diminished Igλ^+^ B cells (Fig.2F). By contrast, the VRC26UCA KI mouse contained higher fractions of Igλ^+^ B cells than F1 mouse (Fig.2G and 2H). Within the Igλ^+^ population in a 7-week old mouse, some B cells expressed abnormally low levels of Igλ (Fig.2G). These Igλ^lo^ B cells likely corresponded to those with low surface IgM^a^ (Fig.2C). In concert with the loss of IgM^a^ B cells in older mice, Igλ^lo^ B cells also disappeared (Fig.2H). Again, the KI mouse with VRC26UCA HC alone had a similar phenotype as the HC/LC mouse, but with a more prominent Igλ^lo^ population (Figure 2M), correlating with larger numbers of IgM^a,lo^ B cells (Fig.2I). The light chain loci were unmodified in the VRC26UCA HC mouse. The increase in Igλ^+^ B cells, and the distinct IgM^a, lo^Igλ^lo^ population, may reflect preferential pairing of VRC26UCA HC with the mouse λ light chain. Expression of VRC26UCA LC also caused a noticeable increase in Igλ^+^ B cells, presumably due to deletion of the KI LC via receptor editing in some B cells (Fig.2O and 2P).

These analyses showed strong negative selection against VRC26UCA, primarily due to its heavy chain. V_H_ replacement can delete rearranged V exons in IgH locus (17, 46). Most V_H_ segments contain a cryptic recombination signal sequence (RSS), which can undergo V(D)J recombination with upstream canonical RSSs to delete the KI V portion of the V(D)J exon and replace it with and upstream V sequence (17, 46), To test whether V_H_ replacement accounted for the loss of VRC26UCA HC expression, we generated hybridomas from splenic B cells of the constitutive mouse model. Consistent with the FACS analysis, the majority of hybridomas secreted IgM^b^ allotype antibodies (Fig.S3A and S3B). Based on PCR analysis, all the examined hybridomas, including those expressing the IgM^a^ allotype antibody, lost the region upstream of the cryptic RSS in VRC26UCA HC (Fig.S3C and S3D). Using a Digestion Circularization-PCR (DC-PCR) method, we identified the sequences that substituted for VRC26UCA HC in some hybridomas (Fig.S3E-S3H, Table S1). These recombination products were typical V_H_ replacement events, in which the cryptic RSS in VRC26UCA HC mediated V(D)J recombination with mouse V_H_ and/or mouse D segments. In hybridomas producing IgM^b^ allotype antibodies, the V_H_ replacement products were nonproductive, including D to VRC26UCA HC rearrangements, out-of-frame V_H_-D-VRC26UCA HC rearrangements, or in-frame V_H_-D-VRC26UCA HC rearrangements with stop codons (Table S1). One IgG1^b^ hybridoma contained an in-frame V_H_-VRC26UCA HC recombination event. It is unclear why this HC was not expressed as an IgM^a^ allotype antibody; one possibility is that the V_H_ replacement product did not fold properly. In hybridomas producing IgM^a^ allotype antibodies, V_H_-D-VRC26UCA HC recombination events were in-frame (Table S1). Thus, in all examined cases, V_H_ replacement eliminated the VRC26UCA HC.

Receptor editing can delete rearranged VJ exons in the Igκ locus, leading to the rearrangement of the other Igκ allele or the Igλ locus (19–22). This process likely accounted for the increase in Igλ^+^ B cells in VRC26UCA KI mouse. Deletion of the VRC26UCA LC via receptor editing could also result in the expression of the other Igκ allele. Without an allotypic marker to distinguish the two Igκ alleles, we could not assess the frequency of such events. Thus, increase in Igλ^+^ B cells likely did not reflect the full extent of VRC26UCA LC deletion. As judged by B cell number (Fig.S2I and S2K), expression of VRC26UCA LC had less of an impact on B cell development than expression of the VRC26UCA HC. Nonetheless, combined expression of both VRC26UCA HC and LC exacerbated B cell deletion.

### B cell Developmental block in the constitutive expression model of VRC26UCA

The analysis above showed that VRC26UCA expression essentially blocked B cell maturation. To define the stage of developmental arrest, we analyzed B cell progenitors in the bone marrow. Based on B220, CD19 and IgM surface markers, we broadly separated bone marrow B cells from the wild-type F1 mouse into three major populations: pre-pro-B cells (B220^+^CD19^−^IgM^−^; population I), pro-B cells and pre-B cells (B220^+^CD19^+^IgM^−^; population II), immature and mature B cells (B220^+^CD19^+^IgM^+^; population III) (Fig.3A; Fig.S4) (47). The bone marrow from B1-8/3-83 KI mouse had minimal numbers of pro-B cells and pre-B cells (population II, Fig.3B). The phenotype is likely attributable to the pre-assembled V(D)J exons in the KI mouse, which obviate the need for V(D)J recombination to assemble HC and LC V(D)J exons in pro-B cells and pre-B cells. The VRC26UCA KI mouse had a different bone marrow B cell pattern from either the F1 mouse or B1-8/3-83 KI mouse (compare Fig.3C with Fig.3A and 3B). Relative to the B1-8/3-3 KI mouse, the VRC26UCA KI mouse had more CD19^+^IgM^−^ pro-B and pre-B cells (compare population II in Fig.3B and 3C). In addition, the bone marrow of VRC26UCA KI mouse contained a prominent B cell population with intermediate levels of CD19 (CD19^int^) and low levels of surface IgM (IgM^lo^) (population IV in Fig.3C). These B cells were IgM^a+^ (Fig.3I), the allotype of VRC26UCA HC, and CD93^hi^, a marker for immature B cells (Fig.3K). A minor population of CD19^hi^IgM^+^ immature and mature B cells was discernable (population III in Fig.3C), but these B cells expressed IgM^b^ mouse antibodies (Fig.3J and 3K). The accumulation, and apparent developmental arrest, of the IgM^a+^CD93^hi^ B cells suggests that maturation of VRC26UCA B cells was impeded at the immature B cell stage. Like splenic B cells, bone marrow B cells from VRC26UCA HC-only mouse exhibited a similar, but less severe, phenotype as those expressing HC and LC KI alleles (compare Fig.3C with 3D, Fig.3I-3K with 3L-3O. The bone marrow B cell profile of VRC26UCA LC alone mouse was largely normal (Fig.3E).

**Fig. 3.**
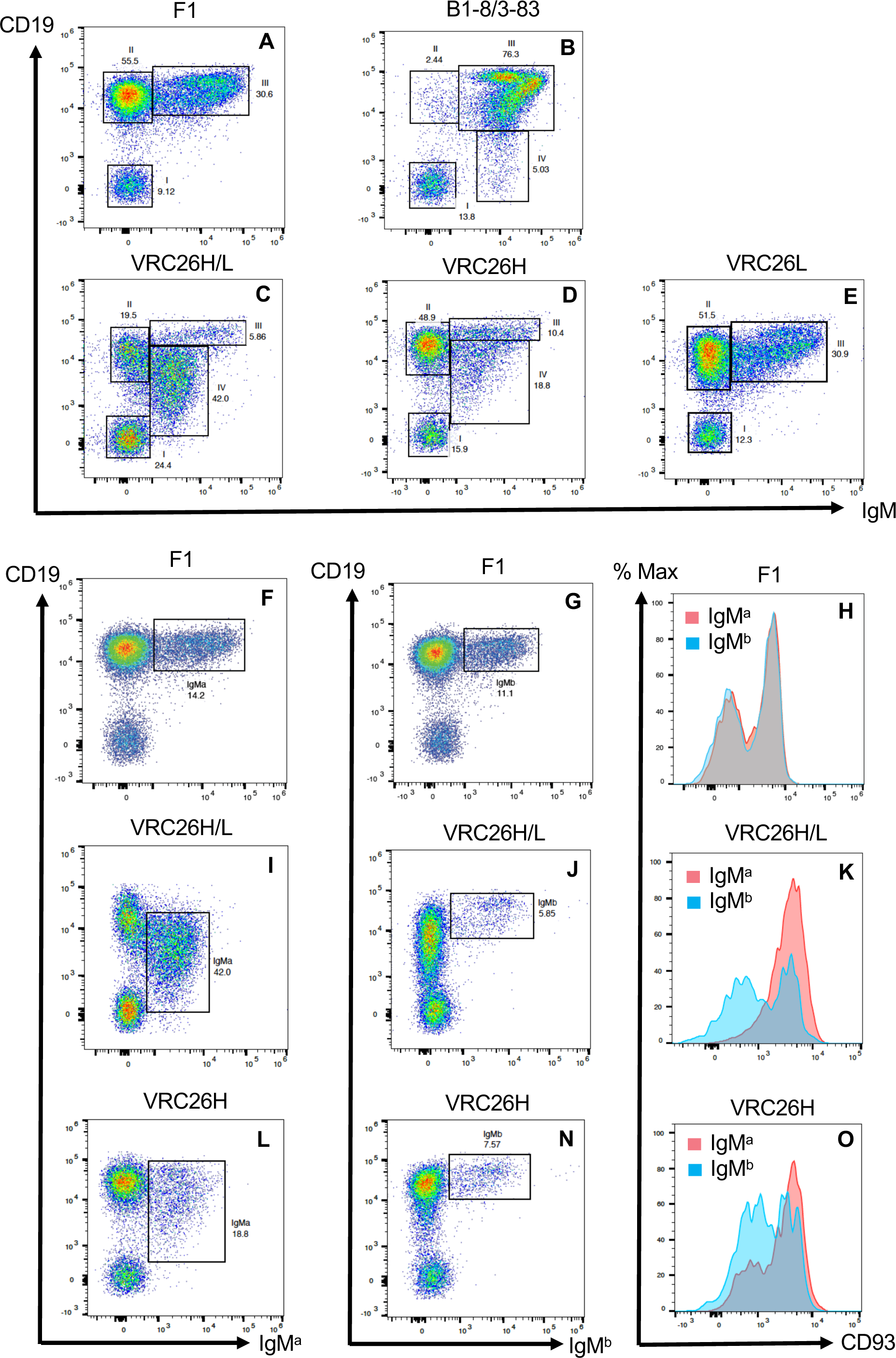
FACS analysis of bone marrow B cells in the constitutive expression model of VRC26UCA. All the plots were gated on B cells. The gating strategy was illustrated in Fig.S4. The plots were labeled in the same manner as those in Fig.2; bone marrow in this figure and spleen in Figure 2 for each genotype were isolated from the same mouse. **(A-E)** These FACS plots show staining patterns of CD19 versus IgM. **(F, I, L)** These FACS plots show staining patterns of CD19 versus IgM^a^. **(G, J, N)** These FACS plots show staining patterns of CD19 versus IgM^b^. **(H, K, O)** These plots show overlays of histograms of CD93 expression from IgM^a^ and IgM^b^ B cells. These two populations differ in size in VRC26UCA KI mice; to compensate for this disparity, the y-axis was normalized to mode, and the highest peak in each distribution is set to 100%. Due to smoothening of the curve, the highest peak may not be visible in some plots.

Some bnAbs are poly-reactive, and expression of such antibodies can trigger tolerance control mechanisms (11–16). This mechanism could underlie the developmental arrest of immature B cells in VRC26UCA KI mouse. To test this possibility, we assessed the cross-reactivity of VRC26UCA to mouse bone marrow and splenocytes. We prefer this assay to the conventional method of antibody staining of HEp2 cells (13), which originated from human epithelial cells and does not represent relevant antigens for B cells in KI mice. For comparison, we used the driver mouse antibody, which was used to support B cell maturation in the conditional expression system. Relative to this control, VRC26UCA exhibited stronger cross-reactivity with B cells from both spleen and bone marrow (Fig.4A and 4C), intermediate binding to non-B cells in bone marrow and non-B/T cells in spleen (Fig.4B and 4E), and no detectable interaction with splenic T cells and red blood cells (Fig.4D and 4F) (See Fig.S5 for definition of these cell populations). This assay did not sample all antigens, such as soluble factors, in bone marrow and in spleen, and could not define the cross-reactive antigens. Despite these limitations, the experiment did reveal the cross-reactive nature of VRC26UCA, providing a potential basis for the developmental arrest at the immature B cell stage in VRC26UCA KI mice.

**Fig. 4.**
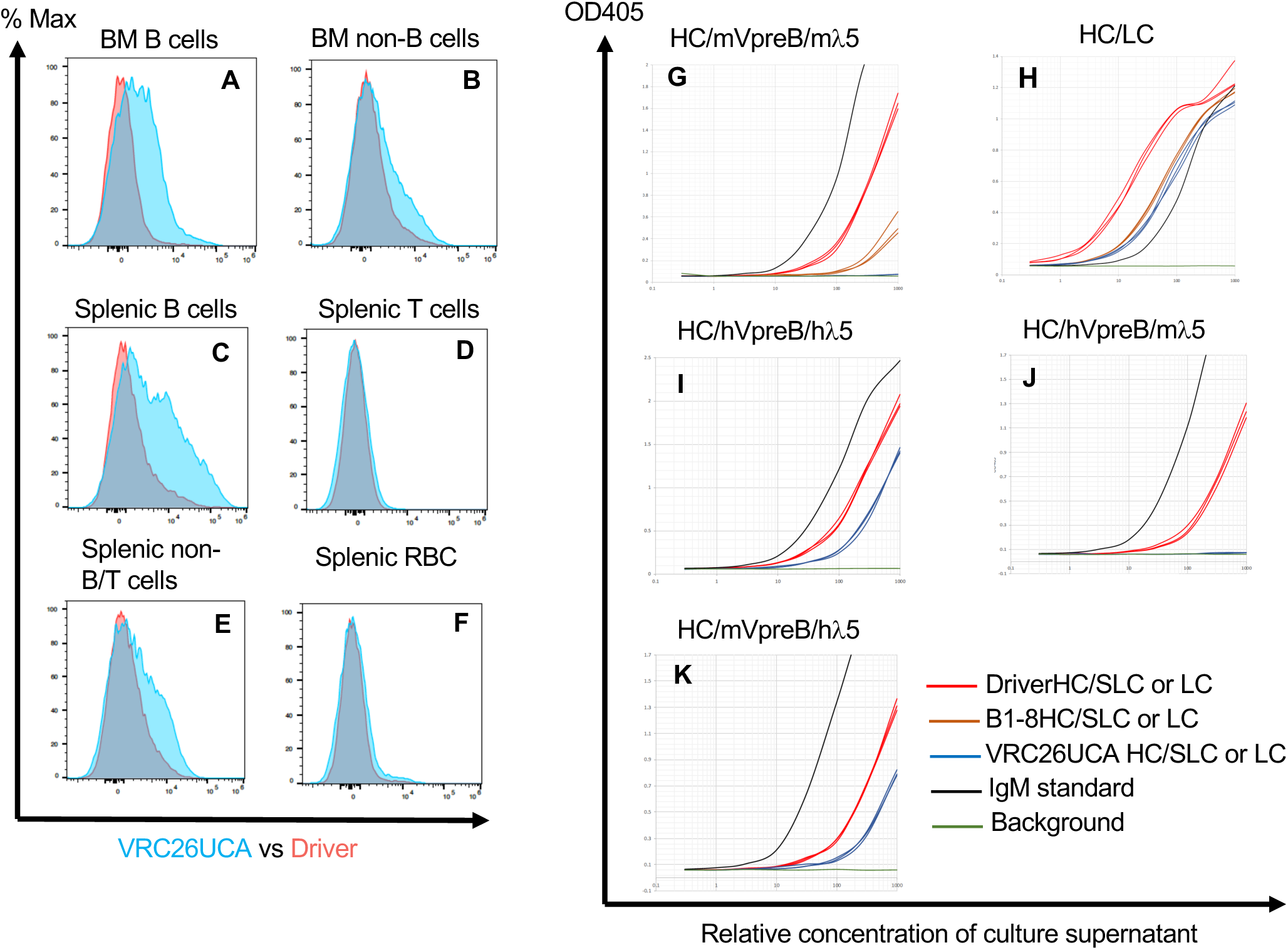
In vitro studies on the potential causes of B cell developmental block. **(A-F)** FACS analysis of cross-reactivity of VRC26UCA antibody. In this experiment, splenocytes or bone marrow cells were stained with VRC26UCA antibody or driver antibody that have been conjugated to fluorophore AF647. As indicated above each plot, binding activity were assessed on B cells and non-B cells from bone marrow (BM) or B cells, T cells, non-B/T cells and red blood cells (RBCs) from spleen. The gating strategy was illustrated in Fig.S5. The binding activity was displayed in histograms, and the plots for VRC26UCA and driver antibodies were overlaid for comparison. The y-axis of the histogram was normalized to mode, as in **Fig.3H, K and O. (G-K)** ELISA analysis of pairing efficiencies of antibody HC with SLC. **(G)** This plot shows the ELISA measurement of the levels of μHC secreted in association with mouse SLC composed of mouse VpreB (mVpreB) and mouse λ5 (mλ5). Each line represents the titration curve of supernatant from one transfection experiment. Each transfection was done in triplicates. Titration curves for different HCs were distinguished by color: driver HC, red; B1-8HC, brown; VRC26UCA HC, blue. Each plot includes a titration curve of IgM standard (black) as a reference; green curve represents background. The x-axis of the plots represents relative concentration of the culture supernatant, and the highest concentration corresponds to undiluted supernatant. The y-axis displays OD405, which correlates with antibody concentration, as measured by μ heavy chain. Plots (H-K) were labelled in the same manner as this plot, except that B1-8HC were not included in the analyses in **I-K. (H)** Levels of μHC secreted in association with LC. **(I)** Levels of μHC secreted in association with human SLC composed of human VpreB (hVpreB) and human λ5 (hλ5). **(J)** Levels of μHC secreted in association with hVpreB and mλ5. **(K)** Levels of μHC secreted in association with mVpreB and hλ5.

In light of the largely empty pro-B and pre-B compartment in B1-8/3-82 KI mouse (population II in Fig.3B), the pre-assembled V(D)J exons of B1-8/3-83 antibody appeared to speed B cell development through these stages. By comparison, the presence of a sizable population of pro-B cells and pre-B cells in VRC26UCA KI mouse (population II in Fig.3C) suggests that, relative to B1-8/3-83, VRC26UCA was less effective in this respect. Furthermore, the pro-B and pre-B cells in VRC26UCA KI mouse eventually developed into IgM^b^ B cells (Fig.3J), which have deleted the VRC26UCA HC via V_H_ replacement and rearranged the other IgH^b^ allele. The two phenotypes, failure to speed-up pro-B/pre-B cell maturation and to suppress V(D)J recombination activity, may be coupled. Both processes depend on the association of nascent HC with surrogate light chain (SLC) to form pre-B cell receptor (48, 49), which promotes the transition from pro-B cells to pre-B cells and terminates IgH rearrangement (50–55); poor pairing of VRC26UCA HC with SLC may lead to defects in both processes. To test this hypothesis, we co-transfected VRC26UCA HC with SLC components, VpreB (56) and λ5 (57), into 293T cells. If the HC does not pair with SLC, the HC should not fold properly for secretion. Thus, the level of secreted HC, presumably in association with SLC, should correlate with their pairing efficiency. For comparison, we performed the same experiment with two mouse HCs: the driver HC and B1-8 HC (Fig.4G and Fig.S6). In support of our prediction, co-transfection of VRC26UCA HC with mouse SLC yielded no detectable HC secretion, in contrast to the results with driver HC and B1-8 HC (Fig.4G). The pairing defect was specific for mouse SLC, as co-transfection of VRC26UCA HC and LC led to robust antibody secretion (Fig.4H). Furthermore, co-transfection of VRC26UCA HC with human SLC also improved antibody secretion (Fig.4I), and human λ5 accounted primarily for the stimulatory effect (Fig.4J and 4K).

### Conditional expression strategy to overcome B cell developmental block

The second goal of this study was to test whether a conditional expression strategy could overcome impediment to B cell development associated with constitutive expression of VRC26UCA. Toward this goal, we generated a conditional expression model of VRC26UCA (Fig.1B). In contrast to the constitutive expression model, where the majority of splenic B cells expressed mouse IgM^b^, most of the splenic B cells in the conditional expression model expressed IgM^a^ (Fig.5B). This result indicated that the driver HC, which was expressed in B cell progenitors, effectively excluded the other IgH^b^ allele and supported B cell maturation. Nonetheless, relative to control IgH^a^ 129Sv mouse, both IgM^a^ and IgD expression on B cells from the conditional expression model were below normal (Fig.5A-F). With respect to Ig light chain expression, the frequency of Igλ^+^ B cells, including a distinct Igλ^lo^ population, was increased (Fig.5G-I). On the other hand, B cells from the conditional expression model of VRC26UCA expressed normal levels of major B cell surface markers, except for a minor increase in class II MHC (I-A^b^) (Fig.S7). Low surface Ig is characteristic of anergic B cells, likely because persistent stimulation by self-antigens down-modulates B cell receptor (24, 25). The poly-reactivity of VRC26UCA (Fig.4A and 4C) may be a contributing factor to the low surface Ig phenotype.

**Fig. 5.**
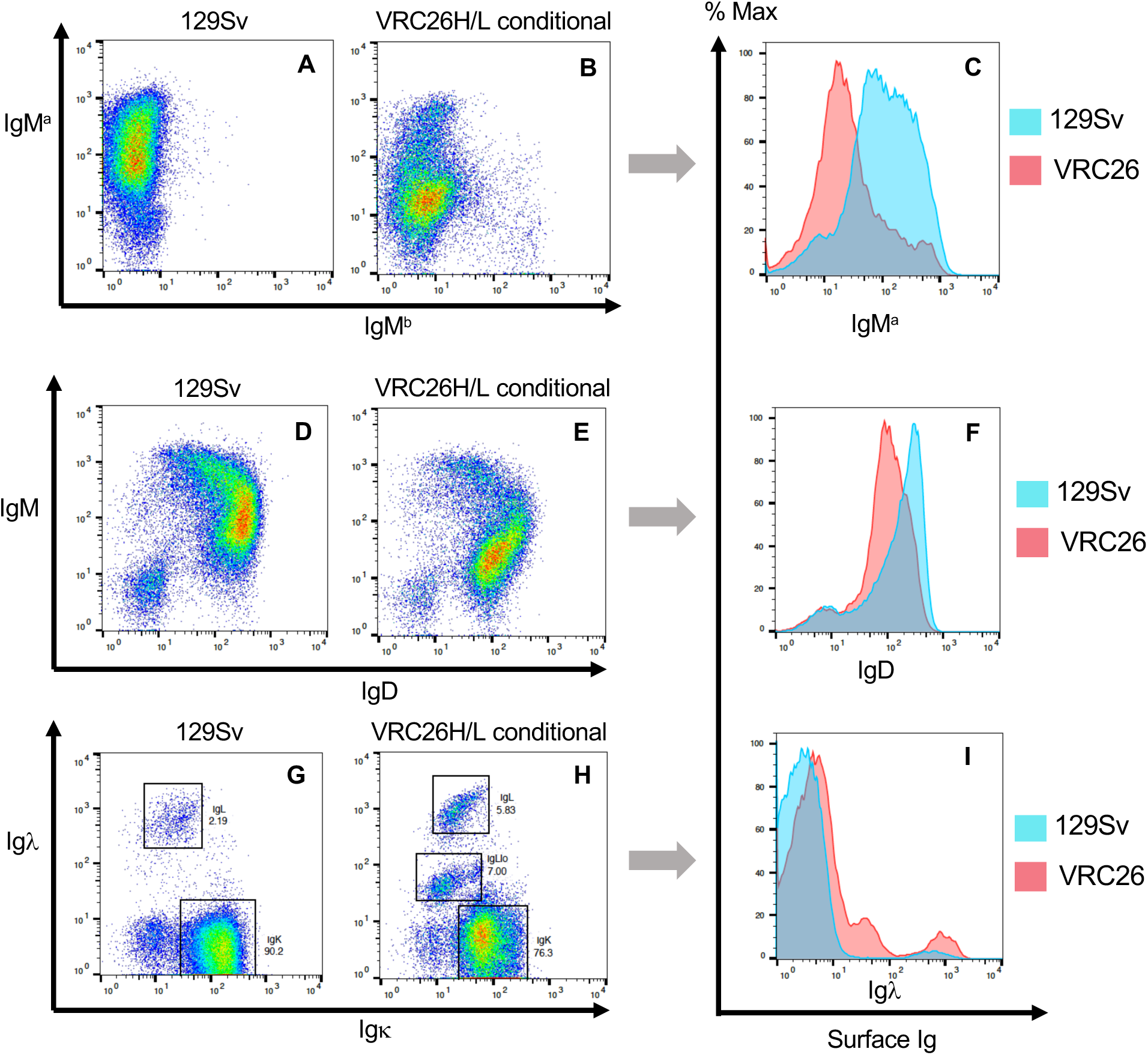
FACS analysis of splenic B cells from conditional mouse model of VRC26UCA. All the plots were gated on B cells, as illustrated in Fig.S2. The staining antibodies were indicated next to the axis. The genotypes of the mice were indicated above the plots: 129Sv, wild-type 129/Sv mouse; VRC26UCAH/L conditional, conditional expression model of VRC26UCA HC and LC. **(A-B)** These plots show the distribution of IgM^a^ versus IgMb expressing B cells. **(C)** The histogram is an overlay of IgM^a^ expression between 129Sv **(A)** and conditional model of VRC26UCA **(B)**. The y-axis of the histogram represents relative cell numbers, and the peak of each histogram is set to 100% (modal scale). **(D-I)** These plots compare the expression patterns of IgM/IgD and Igκ/Igλ between control 129Sv mouse and conditional expression model of VRC26UCA. The plots were labeled as **(A-C)**.

Conditional expression of VRC26UCA depends on deletion of the driver V exon by CD21-cre (Fig.1C). Due to partial deletion, the IgM^a^ B cell population consisted of a mixture of VRC26UCA and driver expressing B cells. To determine the ratio of these two types of B cells, we used the same DC-PCR method as for V_H_ replacement analysis (Fig.S8). Because of its long CDR H3, the DC-PCR product for VRC26UCA HC is longer than that of the driver HC (Fig.6 and Fig.S8B). Control experiments confirmed unbiased amplification of VRC26UCA HC and driver HC with this DC-PCR method (Fig.S8B). Thus, the ratio of the two DC-PCR products should reflect the fraction of B cells that express the two antibodies. Strictly speaking, the method detects variable region exons that are positioned for expression at the DNA level, but not the actual expression of the variable region at RNA or protein level. Since B cells are dependent on functional B cell receptors for survival (58), it is reasonable to assume that the V(D)J exon, which is proximal to the constant regions, is expressed. Since CD21-cre acts primarily in mature B cells, VRC26UCA expression, as a consequence of driver deletion, should happen predominantly in mature B cells. We sorted splenic B cells into IgM^+^IgD^hi^ and IgM^+^IgD^lo^ populations (Fig.6A), which were enriched for mature and immature B cells, respectively. Based on the DC-PCR assay, about 30% of mature splenic B cells expressed VRC26UCA, and consistent with the expression pattern of CD21-cre, few immature B cells expressed VRC26UCA (Fig.6B). VRC26UCA expression was further enriched among B cells with lower IgM expression (Fig.6C-D), presumably owing to poly-reactivity of this antibody.

**Fig. 6.**
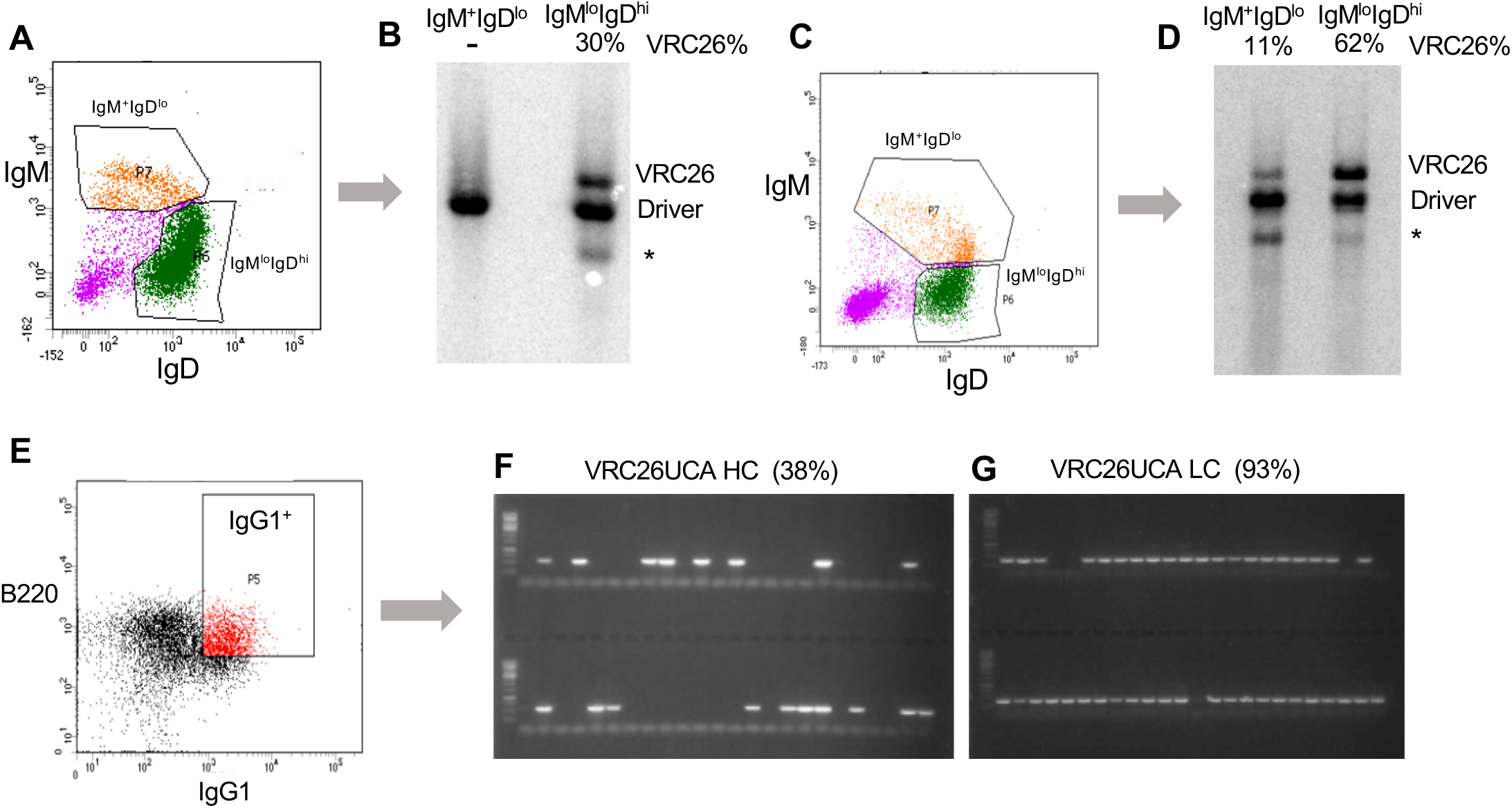
Analysis of the fraction of B cells expressing VRC26UCA in the conditional expression model. **(A-D)** DC-PCR analysis of the ratio of splenic B cells expressing driver or VRC26UCA. **(A)** The FACS plot shows the B cell populations that were sorted for DC-PCR analysis: P6, IgM^+^IgD^hi^; P7, IgM^+^IgD^lo^. The plot was gated on B220^+^ B cells. **(B)** DC-PCR analysis was performed on sorted B cells from **(A)**. The PCR products were run on agarose gel, detected with Southern hybridization and quantified with PhorphorImager. The identity of the PCR products was labeled to the right of the autoradiogram; an unknown PCR product was indicated with *. At the top of the autoradiogram, the B cell population analyzed in this assay was indicated; VRC26% =VRC26/(VRC26+driver). Panels **C** and **D** show a similar experiment as panels **A** and **B**, except that the P6 gate was shifted toward lower IgM levels. **(E-G)** Single-cell RT-PCR assay to determine the frequency of activated IgG1+ splenic B cells expressing VRC26UCA in the conditional expression model. **(E)** Splenic B cells from the conditional expression model were activated in vitro with anti-CD40 antibody plus IL4. After 3 days of activation, B220+IgG1+ activated B cells (P5) were sorted as single-cells into 96-well plate. **(F)** VRC26UCA HC or **(G)** LC transcripts were amplified from the sorted B cells. Each lane corresponds to PCR reaction of one cell. Only part of the gel image was shown. The percentage of VRC26UCA HC or LC positive B cells out of the total sorted B cells, 95 in this experiment, was indicated at the top of the gel image.

The low surface Ig phenotype of VRC26UCA B cells raised concerns about their function. As a preliminary assessment of their functional status, we stimulated splenic B cells from the VRC26UCA mice with anti-CD40 antibody plus IL-4, which activates B cells to undergo IgH class switching to IgG1 and IgE. We sorted IgG1^+^ B cells and performed single-cell RT-PCR (Fig.6E). About 38% B cells expressed VRC26UCA HC (Fig.6F), and 90% of B cells expressed VRC26UCA LC (Fig.6G). This result showed that VRC26UCA expressing B cells can at least be activated *in vitro*. Furthermore, the analysis confirmed the expression of VRC26UCA at the RNA level in a substantial fraction of splenic B cells, in corroboration with the DC-PCR analysis. As a validation of the conditional expression strategy, we also confirmed that VRC26UCA expression was dependent on driver deletion (Fig.S9).

## Discussion

We show that constitutive expression of VRC26UCA caused a B cell developmental arrest, primarily at the immature B cell stage. These immature B cells had atypical patterns of surface markers: intermediate CD19 and low IgM (Fig.3C). Bone marrow from wild-type mouse does not contain substantial numbers of B cells with this surface phenotype (Fig.3A). Such B cells have also not been reported in other bnAb KI mice, but bone marrow B cells were not examined with the combination of CD19 and IgM markers in those studies (5–10). Since a similar CD19^int^ B cell population was also detectable in B1-8/3-83 mouse (Fig.3B), this type of B cell may be present, in varying numbers, in other KI mice. The origin of this B cell population is unclear. One possibility is that these B cells develop conventionally through pro-B cells and pre-B cells, but down-modulate both CD19 and IgM at the immature B cell stage, presumably due to poly-reactivity of the nascent B cell receptor. However, the presence of this type of B cells in both VRC26UCA and B1-8/3-83 KI mice led us to favor the possibility that the phenomenon may relate more generally to the pre-assembled V(D)J exons. Since expression of pre-assembled V exons does not depend on V(D)J recombination in pro-B cells and pre-B cells, HC and LC expression could potentially begin earlier than usual, during the transition from CD19^−^ pre-pro-B cells to CD19^+^ pro-B cells (59), hence their CD19^int^ phenotype. In this scenario, these B cells essentially skipped the normal pro-B cell and pre-B cell stages. For some antibodies, such as B1-8/3-83, B cells could apparently reach maturity through this pathway. However, in the case of VRC26UCA and perhaps other antibodies, the non-physiological maturation pathway might exacerbate the negative impacts of other factors, such as poly-reactivity. One way to address this developmental problem is to express bnAbs through *de novo* assembly of their V(D)J exons via V(D)J recombination, thereby enforcing the physiological maturation pathway for B cell progenitors. We previously took this approach for expressing precursors for VRC01 class antibodies in mice (60). B cells in this VRC01 model developed normally, and immunization elicited VRC01-lineage antibodies.

Not all B cells expressed surface IgM prematurely, prior to pro-B cell stage, in VRC26UCA KI mouse; the CD19^+^IgM^−^ population (population II in Fig.4C) in VRC26UCA KI mouse may consist of such B cells. This population was more numerous in the VRC26UCA KI mouse than in B1-8/3-83 KI mouse. We interpret this difference as a reflection of inefficiency of VRC26UCA, even when expressed as a pre-assembled V exon, in driving B cell maturation through pro-B and pre-B cell stages. The abundant V_H_ replacement events and rearrangement of the other mouse IgH allele in the VRC26UCA KI mouse also suggests that the VRC26UCA HC failed to effectively suppress V(D)J recombination and to expedite B cell maturation through the pro-B cell stage. These observations lead us to propose that poor pairing of VRC26UCA HC with mouse SLC may underlie these phenomena, and our *in vitro* experiments support this hypothesis. The basis for the pairing defect is unclear, but the extraordinarily long CDR H3 of VRC26UCA HC could be relevant. It has been shown that some antibodies with long CDR H3s are under negative selection during B cell maturation (36), and SLC could play a role in this selection. The non-Ig domains of VpreB and λ5 form the equivalent of CDR L3 and interact with CDR H3 of the HC (61). It is possible that long CDR H3, at least in some cases, structurally clashes with the non-Ig domain of the SLC. The 37 amino acid CDR H3 of VRC26UCA is far above the average CDR H3 length of mouse antibodies, and the mouse SLC may not be adapted to accommodating such extraordinarily long CDR H3s. By comparison, antibodies with long CDR H3s are more common in human than in mouse repertoires (34, 35, 62). The difference may explain more efficient pairing of VRC26UCA HC with human SLC than with mouse SLC. To test the physiological relevance of these *in vitro* observations, an obvious experiment is to swap mouse SLC with human counterpart and determine whether human SLC facilitates B cell maturation in this and potentially other bnAb mouse models. Besides testing our hypothesis outlined above, the measure could also alleviate B cell developmental defects in other bnAb KI mice (5–10).

One implication this study is that multiple factors may contribute to B cell defects in bnAb KI mice. Some of these factors, such as a non-physiological B cell developmental pathway and mouse SLC pairing defect are specific to the KI mouse system and may not be relevant to human vaccination. Eliminating these non-physiological factors would not only increase target B cells for immunogens, but also reveal physiologically relevant hurdles for bnAb induction, and the information could facilitate the development of intervention strategies. The conditional expression approach is one step in this direction. The method bypasses the obstacles at the B cell progenitor stage, including some potentially non-physiological hindrances, and generates mature B cells expressing the target antibody for testing immunogens. In the case of the conditional expression model of VRC26UCA, the mature B cells express abnormally low levels of surface Ig and may be functionally anergic. Activating such B cells will likely be challenging. On the other hand, if the anergic state reflects *bona fide* peripheral tolerance, the model could be used to develop strategies to revitalize the anergic B cells, for example with multimeric immunogens or potent adjuvants. Given the prevalence of poly-reactivity among bnAbs, such intervention may be necessary for bnAb induction (30, 63). The conditional expression method is obviously not a feasible way to overcome deletion of bnAb precursors in humans. The value of the strategy lies in generating mouse models with sufficient target B cells for testing and optimizing immunization strategies at the preclinical stage. In human vaccination, an effective vaccine most likely will need to activate rare target B cells that have survived tolerance control checkpoints or other restrictions. Vaccine optimization in the model system we describe here, or in related mouse model systems, may increase the chance of success in overcoming such potential hurdles.

## Materials and Methods

### Generation of constitutive and conditional expression model of VRC26UCA

The conditional expression system requires a CD21-cre transgene. To expedite the setup of the conditional expression system, we derived an ES cell line from CD21-cre transgenic mice (39). The genetic background of the original CD21-cre transgenic mouse line is C57BL/6. As described in the text, we planned to utilize IgH allotypic markers to differentiate B cells expressing KI VRC26UCA antibody versus mouse antibodies. Toward this end, we crossed the CD21-cre mice with 129Sv mice and derived an ES cell line from 3.5-day embryo. In this CD21-cre ES cell line, one IgH allele is a allotype from 129/Sv mouse strain, whereas the other IgH allele is b allotype from C57BL/6 mouse strain. We used this CD21-cre ES cell line for incorporating VRC26UCA HC and LC expression constructs.

The organization of the integration construct for VRC26UCA HC is illustrated in Fig.S1A; the diagram also applies to the LC KI construct. The homology arms of the integration constructs were amplified with high-fidelity PCR from the genomic DNA of an 129Sv ES cell line. Linearized construct was electroporated into the CD21-cre ES cell line described above. Clones with stable integration of the construct were selected with G418. These clones were screened by Southern blotting. Correct clones were transduced with Adeno-cre virus. As illustrated in Fig.S1A, partial recombination with the three loxP sites in the construct gave rise to ES clones for either constitutive expression or conditional expression of VRC26UCA. LoxP recombination pattern was determined by Southern blotting. From this screening, we selected ES clones for constitutive or conditional expression of VRC26UCA. All clones for generating mouse models were verified for genotype (Fig.S1B and S1C) and karyotype (96% ES cells with 40 chromosomes for the conditional expression model; 90% ES cells with 40 chromosomes for the constitutive expression model).

To generate the KI mice, the ES cells were injected into Rag2 deficient blastocysts. Since Rag2 is essential to V(D)J recombination, B and T cells in the chimeric mice are derived from the injected ES cells, but not from the blastocysts. Thus, the chimeric mice can be used directly for analysis of KI antibody expression and B cell development. We refer to this technique as Rag2 deficient blastocyst complementation (RDBC) (64). For long-term studies, we also bred the chimeric mice for germline transmission. In the present study, all analyses of the constitutive expression model were performed with germline KI mice that resulted from cross between VRC26UCA KI mice and C57BL/6 mice. These mice are of mixed 129Sv and C57BL/6 genetic background; the KI VRC26UCA HC was expressed as IgH^a^, whereas mouse antibodies were IgH^b^. The conditional expression model also gave germline transmission. However, presumably due to leaky expression of CD21-cre in germ cells, the driver V gene flanked with loxP sites (floxed Driver) tended to be deleted in a substantial fraction of progenies during breeding. This issue compounded the difficulty of obtaining mice with genetic modifications on three chromosomes (HC KI, LC KI, CD21-cre) by breeding. For this reason, we used RDBC chimeric mice for the analysis of the conditional expression model.

### Characterization of splenic and bone marrow B cells in mouse models

For splenic B cell analysis, spleen was dissociated mechanically into cell suspension. Red blood cells were eliminated by Red Blood Cell lysing buffer (Sigma R7757). To generate the data in Fig.2, Fig.S2, and Fig.5, the splenocytes were stained with the following antibodies: PE-Cy5 anti-B220 (eBioscience 15-0452-83), PE anti-Thy1.2 (PharMingen 553006), PE anti-IgM (eBioscience 12-5790-83), PE anti-IgM^a^ (PharMingen 553517), PE anti-Igλ (Biolegend 407308), FITC anti-IgD (BD PharMingen 553439), FITC anti-IgM^b^ (PharMingen 553520), FITC anti-Igκ (SouthernBiotech 1050-02). The stained cells were analyzed on BD FACS Calibur, and FACS plots were generated with FlowJo10 software. For analysis of various B cell marker expression in Fig.S7, splenic B cells were stained with the following additional antibodies: PerCP/Cy5.5 anti-B220 (eBioscience 45-0452-82), APC anti-IgM (eBioscience 17-5790-82), APC anti-IgD (Biolegend 405713), PE anti-CD93 (eBiosceince 12-5892-82), PE anti-CD22 (SouthernBiotech 1580-09), PE anti-I-A^b^ (SouthernBiotech 1896-09), PE anti-CD86 (SouthernBiotech 1735-09), PE anti-CD40 (eBioscience 12-0401-81), PE anti-CD19 (BD Pharmingen 557399), PE anti-CD23 (BD Pharmingen 553139), FITC anti-CD21 (BD Pharmingen 553818). The stained cells were analyzed on the Attune NxT from Invitrogen, and the data were analyzed with FlowJo 10 software.

For bone marrow B cell analysis, bone marrow cells were flushed out from femur and tibia with syringe. Red blood cells were eliminated with Red Blood Cell lysing buffer (Sigma R7757). In the experiments shown in Fig.3A-3E and Fig.S4, bone marrow cells were stained with the following antibodies: FITC anti-B220 (eBioscience 11-0452-85), APC anti-CD19 (SouthernBiotech 2018-01), PE anti-IgM (eBioscience 12-5790-83), PE-Cy7 anti-CD43 (BD Pharmingen 562866), Sytox blue (S34857). Since CD43 expression was not discussed in this study, CD43 staining was not shown in the figures. For the experiments shown in Fig.3F-3O, bone marrow cells were stained with FITC-anti-B220 (eBioscience 11-0452-85), APC-anti-CD19 (SouthernBiotech 2018-01), PerCP Cy5.5 anti-CD93 (eBioscience 45-5892-82), PE-anti-IgM^a^ (PharMingen 553517) or PE anti-B220 (eBioscience 12-0452-82), APC anti-CD19 (SouthernBiotech 2018-01), PerCP Cy5.5 anti-CD93 (eBioscience 45-5892-82), FITC anti-IgM^b^ (PharMingen 553520). Flow cytometry was performed on Attune NxT from InVitrogen. The data were analyzed with FlowJo10 software.

### Analysis of V_H_ replacement

Splenocytes were isolated from a constitutive expression model of VRC26UCA and stimulated with anti-CD40 antibody (Invitrogen 16-0402-86) plus IL4 (65). After 3 days of stimulation, activated B cells were fused with NS1 plasmacytoma cells with the PEG method (66). Hybridoma clones were selected with HAT. Supernatants from the hybridoma clones were screened via ELISA to determine the isotype and allotype of secreted antibodies. Simulation with anti-CD40 plus IL4 induces class switching to IgG1 and IgE (65). Unswitched B cells continue to express IgM. As a result, most of the hybridoma clones secreted IgM, IgG1 or IgE. Some supernatants contained more than one antibody isotype; these samples may be derived from mixed hybridoma clones and were not used for further analysis. Hybridoma clones producing only IgM or IgG1 were used for the experiments in Fig.S3. The supernatants from these hybridomas were further screened with allotype-specific antibodies. For IgM^+^ hybridomas, their supernatants were screened with anti-IgM^a^ and anti-IgM^b^ antibodies. Hybridoma clones expressing only IgM^a^ or IgM^b^ were chose for further analysis. For IgG1^+^ hybridomas, their supernatants were screened with anti-IgG1^a^ antibody only; the anti-IgG1^b^ antibody works poorly in ELISA assays. IgG1^a-^/IgG1^+^ hybridomas were considered as IgG1^b+^. Without direct identification of IgG1^b^ hybridomas, some IgG1^a+^ hybridomas could be mixed with IgG1^b+^ hybridomas, causing over-estimation of the frequency of IgG1^a^ hybridomas.

In Fig.S3C, genomic DNAs from IgM^+^ hybridomas were amplified with primers that anneal to sequences downstream of the cryptic RSS. In Fig.S3D, the same DNA samples from Fig.SC were amplified with primers that anneal to sequences upstream and downstream of the cryptic RSS, respectively. To amplify V_H_ replacement products with DC-PCR, genomic DNA from hybridomas were digested with NcoI and XbaI or PstI and XbaI. The digested DNA was self-ligated into circles with adaptor primers that have cohesive ends for NcoI and XbaI or PstI and XbaI. The ligated product was subject to PCR amplification with primers e and f (Fig.S3E). The PCR product was sequenced. The identities of the mouse D or V_H_ segments joined to VRC26UCA HC, as shown in Fig.S3F, S3G, and S3H and Table S1, were determined with Blast or Ig Blast.

### Assessment of cross-reactivity of VRC26UCA antibody

To produce recombinant VRC26UCA antibody and the driver antibody, the cDNAs for the heavy and light chains of these antibodies were cloned into pcDNA3 expression vector. The constant regions for the heavy and light chains were mouse μ and κ isotypes, respectively. A 6xHis tag was appended to the C-terminus of the heavy chain to facilitate purification. The expression constructs were transfected into Expi293 cells, using ExpiFectamine (Gibco A14524). The supernatant of the culture was harvested 5-6 days after transfection. Antibodies were purified from the supernatant with HisTrap column on FPLC. The purified antibody was conjugated to AlexaFuro647 with microscale protein labeling kit (Invitrogen A30009). For the cross-reactivity assay, bone marrow cells or splenocytes were stained with the following antibodies: FITC anti-B220, PE anti-Thy1.2, Biotin Ter119 (Biolegend 116203), BV605 Streptavidin (Biolegend 405229), PE-Cy7 anti-TNP (Biolegend 401627), AF647 VRC26UCA or AF647 Driver. Stained splenocytes or bone marrow cells were analyzed on an Attune flow cytometer from Invitrogen. FACS plots were generated with FlowJo10 software.

### Analysis of heavy chain pairing with surrogate light chain

The heavy chain, light chain, surrogate light chain cDNAs were cloned into pcDNA3 expression vector. The expression constructs were transfected into 293T cells, using PEI method (67). To control for transfection efficiency, a pMaxGFP expression construct (Lonza) was co-transfected with antibody expression constructs. The ratio of heavy chain, light chain and GFP expression constructs was: 50:50:5; the ratio of heavy chain, VpreB, lambda 5 and GFP expression constructs was: 33:33:33:5. Two days after transfection, the supernatant of the culture was collected. Antibody concentration in the supernatant was measured with ELISA, using anti-mouse IgM antibody (SouthernBiotech 1021-01) for capture and anti-mouse IgM-alkaline phosphatase antibody (SouthernBiotech 1021-04) for detection. Purified mouse IgM (SouthernBiotech 0101-01) served as standard.

### DC-PCR assay to determine the frequency of splenic B cells expressing VRC26UCA in the conditional model of VRC26UCA

Splenic B cells were sorted into B220^+^IgM^+^IgD^lo^ and IgM^+^IgD^hi^ fractions on BD FACS Aria. Genomic DNA was isolated from the sorted B cells. The genomic DNA was digested with NcoI and EcoRI. The digested DNA was ligated with adaptor that contains compatible cohesive ends with NcoI and EcoRI. In the ligation reaction, low DNA concentration favors intramolecular recircularization. Circularized DNA containing the variable regions of VRC26UCA HC or driver HC share a common region downstream of J_H_. PCR amplification with primers a and b, which anneal to the common sequence, yielded products containing the two variable regions at similar efficiencies. The PCR product was run on agarose gel. The DNA was transferred to Zeta-Probe GT membrane (Bio-Rad 162-0197). The DNA was hybridized to an oligonucleotide probe that is common to the PCR products of both variable regions. The hybridization signal was quantified with PhorphorImager (GE Storm 865).

### Single-cell RT-PCR assay to determine the fraction of B cells expressing VRC26UCA in **activated splenic B cells**

Splenic B cells were isolated from the conditional expression model of VRC26UCA. The B cells were stimulated with anti-CD40 antibody plus IL-4. After three days of stimulation, single IgG1^+^ B cells were sorted into 96-well plate. Reverse transcription of the RNA from the sorted single cells yielded cDNA, which served as template for PCR amplification for VRC26UCA HC and LC. VRC26UCA HC cDNA was amplified with forward primer for the HC variable region and reverse primer for Cγ1. Likewise, VRC26UCA LC cDNA was amplified with forward primer for the LC variable region and Cκ.

### PCR amplification of VRC26UCA HC or driver HC from hybridomas of conditional expression model of VRC26UCA

Hybridomas were generated and screened with the same method as that for the constitutive model, as described above. Fig.S9 shows the analysis of 8 IgG1^+^ hybridomas. As illustrated in the diagram above each gel image, RT-PCR amplification of driver or VRC26UCA HC transcripts were achieved with forward primers specific for each variable region and reverse primer in Cγ1. PCR amplification of the driver HC or VRC26UCA HC directly from hybridoma DNA was achieved with forward primers specific for each variable region and a reverse primer downstream of J_H_.

## Acknowledgements

We thank our colleagues at Duke Human Vaccine Institute, Dr. John Mascola and colleagues at Vaccine Research Center for discussion throughout the study. Peiyi Hwang performed mouse work, including embryo microinjection, for the early stage of the project. Author contributions: MT, BH and FA designed and supervised the project. MT, KM, HC, PW, LR, MD, YC, MK, EC, NM, RJ, AW and DC performed the experiments. MT and FA wrote the manuscript. This work was supported by a Division of AIDS UM1 grant AI100645 for the Center for HIV/AIDS Vaccine Immunology-Immunogen Discovery (CHAVI-ID) to B.F.H and to F.W.A, and by a Division of AIDS UM1 grant AI144371 for the Consortium for HIV/AIDS Vaccine Development (CHAVD) to B.F.H and to F.W.A. FWA is an Investigator of the Howard Hughes Medical Institute.

## Supplementary figure legends

**Fig.S1.** Generation and genetic validation of the constitutive and conditional expression models of VRC26UCA.

**(A)** Illustration of the strategy to generate constitutive and conditional mouse models of VRC26UCA. The KI construct consists of 5’ homology arm (5’HL), V exon for VRC26UCA HC with upstream V_H_ promoter (VRC26), neomycin resistance gene (neo^r^), V exon for driver HC with upstream V_H_ promoter (Driver), 3’ homology arm (3’HL); the position of 3 loxP sites were indicated. To simplify the diagram, two poly(A) sites, between loxP-2 and Driver, were not depicted. Homologous recombination between the two homology arms with corresponding sequences in the J_H_ locus integrated the expression cassette into the J_H_ locus in ES cells. Transduction of the ES clones with limiting amounts of adenovirus-cre led to partial recombination between the three loxP sites. Recombination between loxP-1 and loxP-3 generated the ES clone for the constitutive expression model of VRC26UCA. Recombination between loxP-1 and loxP-2 generated the ES clone for the conditional expression model of VRC26UCA. VRC26UCA LC was integrated into the mouse Jκ locus with a similar strategy. In this case, ES clones for constitutive expression of VRC26UCA LC were chosen to generate the mouse models for both the constitutive and conditional expression models of VRC26UCA, as illustrated in Figure 1. **(B)** Correct integration of the KI constructs was verified by Southern blotting of the ES clones. Integration of the KI constructs altered the pattern of certain restriction digests. The change was revealed by Southern hybridization with probes that are external to the homology arms. The use of external probes ensures that the hybridization reveals restriction digest pattern at the endogenous locus. Two probes, which lie 5’ or 3’ to the homology arms, were used to verify both ends of the integration. The table lists the predicted sizes of the restriction digest fragments that are detectable by each probe. **(C-F)** Southern hybridization analysis of ES clones for constitutive or conditional expression model of VRC26UCA. The probe for each hybridization were indicated below the Southern blot. The genomic DNA was digested with corresponding enzymes for each probe, as listed in the table in **(B)**. The sizes of DNA marker were indicated to the left of each blot. Lanes 1-3 contained genomic DNAs from parental ES cell, ES clone for constitutive expression and ES clone for conditional expression. Based on this analysis, both the constitutive and conditional clones exhibited correct restriction digest pattern.

**Fig.S2.** FACS Analysis of splenic B cells of constitutive expression model of VRC26UCA.

**(A** and **B)** Illustration of the gating strategy for the FACS plots in Fig.2, which were derived from the B cell gate in **B**. Panels **C-K** show the relative ratios of B versus T cells. For this analysis, splenocytes were stained with PE/Cy5 anti-B220 and PE anti-Thy1.2 antibodies. All the FACS plots from **C-K** were gated on lymphocytes, as defined in panel **A**.

**Fig.S3.** Analysis of VH replacement events in the constitutive expression model of VRC26UCA.

**(A-B)** The pie charts summarized the number of IgM^a^, IgM^b^, IgG1^a^, IgG1^b^ hybridomas that were derived from splenic B cells of a constitutive expression model of VRC26UCA. Without a good anti-IgG1^b^ antibody in ELISA assay, the number of IgG1^b^ hybridoma was derived indirectly by subtracting the number of IgG1^a^ hybridomas from total IgG1 hybridomas. The total numbers of IgM or IgG1 hybridomas were shown at the center of the pie charts. The number of IgM^a^, IgM^b^, IgG1^a^ and IgG1^b^ hybridomas were indicated to the right of each pie chart. **(C)** Genomic DNA from 20 hybridomas (lanes 1-20) and liver DNA from a VRC26UCA KI mouse as positive control (lane C) were subject to analysis for V_H_ replacement events. 19 of the hybridomas secreted IgM^b^ antibody; one hybridoma (lane 17) secreted IgM^a^ antibody. The DNA was subject to PCR amplification with a pair of primers downstream of the cryptic RSS in VRC26UCA HC. **(D)** The same experiment as in **(C)**, except that one primer annealed upstream of the cryptic RSS. **(E)** Illustration of the DC-PCR method to clone V_H_ replacement products. **(F)** Diagram of the V(D)J exon of VRC26UCA HC, which includes human VH3-30 gene segment, D3-3 segment and J_H_3 segment; N represents nucleotides between V_H_ and D, D and J_H_ segments. The black triangle in VH3-20 represents the cryptic RSS (cRSS). The sequence surrounding the cryptic RSS was shown below. **(G)** Diagram of V_H_ replacement product involving mouse D segment (mD) and VRC26UCA HC. The blank area between mouse D segment and VRC26UCA HC, in orange, represents N nucleotides. An example of this type of V_H_ replacement product was shown below: mouse D region sequences, in blue, and N nucleotides, in black, were italicized; the D segment was underlined. The remnant of VRC26UCA HC was shown in orange and was aligned with the original sequence in **(F)** for comparison. **(H)** Diagram of V_H_ replacement product involving mouse V_H_ segment, mouse D segment and VRC26UCA HC. This figure was organized in the same manner as **(G)**, except for the addition of mouse V_H_ segment in red.

**Fig.S4.** Illustration of the gating strategy for the analysis shown in Fig.4.

FACS plots **A**-**D** in this figure were based on analysis of bone marrow from F1 mouse shown in Fig.3A, which was derived from the B220^+^ gate in plot **D**. All FACS plots in Fig.4 were based on the same gating strategy.

**Fig.S5.** Illustration of the gating strategy for the analysis of cross-reactivity of VRC26UCA and driver antibody with bone marrow cells and splenocytes in Fig.4.

Staining of bone marrow B cells (Fig.4A) and non-B cells (Fig.4B) was gated on B220^+^ and B220^−^ populations in panel **D**, respectively. Staining of splenic B cells (Fig.4C), T cells (Fig.4D), non-B/T cells (Fig.4E) and red blood cells (Fig.4F) were based on B220^+^Thy1.2^−^, B220^−^, Thy1.2^+^, B220^−^Thy1.2^−^ gates in panel **H** and B220^−^Ter119^+^ gate in **K**.

**Fig.S6.** Control for transfection efficiency in the HC/SLC paring analysis.

In the transfection experiments shown in Fig.6, a GFP expression construct was co-transfected with those for antibodies. The fraction of GFP^+^ cells served as control for transfection efficiency. Plots **A-C** show the FACS analysis of GFP^+^ cells, using one of transfection experiments for driver HC with mVpreB and mλ5, as shown in Fig.4G, as an example. The tables in **D-H** listed the fraction of GFP^+^ cells in each of transfection experiments shown in Fig.6. Based on these data, the transfection efficiencies in different experiments were comparable.

**Fig.S7.** Comparison of the expression levels of various B cell markers on splenic B cells from control C57BL/6 mouse or the conditional expression model of VRC26UCA.

Panels **A** and **B** show FACS plots of IgM/IgD expression of splenic B cells from C57BL/6 mouse or conditional expression model of VRC26UCA; the plots were gated on B220^+^ B cells as illustrated in Fig.S2. IgM^+^IgD^hi^ population, gated in this plot, was further assessed for expression of various B cell markers, displayed in histograms. The histograms from C57BL/6 mouse and the conditional expression model of VRC26UCA were overlaid **(C-J)**. In these histograms, x-axis represents the expression levels of various B cell markers; y-axis represents relative cell number, with the peak of each histogram set at 100%.

**Fig.S8.** Description of the DC-PCR assay to measure the ratio of B cells expressing driverHC or VRC26UCA HC in splenic B cells.

**(A)** In the DC-PCR assay, genomic DNA was digested with NcoI (N) and EcoRI (E). The digested DNA was ligated with an adaptor for NcoI and EcoRI sites into circles. The circular DNA was subject to PCR amplification with primers a and b. The PCR reaction yielded products that contained the variable regions of either driver HC or VRC26UCA HC. The PCR products were run on agarose gel and detected by Southern blotting (Fig.6B **and 6D)**. **(B)** To determine the ratio of driver HC and VRC26UCA HC, the DC-PCR method must amplify the two variable regions equally. To validate this point, genomic DNA was isolated from the liver of a mouse that was heterozygous for constitutive and conditional expression alleles of VRC26UCA. The driver V region in the conditional expression cassette was not deleted in liver cells. In this DNA sample, the template ratio for driver HC and VRC26UCA HC in DC-PCR assay should be 1:1. After DC-PCR, the products were run on an agarose gel: M, 1kb DNA ladder; lane 1, DC-PCR without ligation adaptor, lane 2, DC-PCR with ligation adaptor. Due to its extraordinarily long CDR H3, the PCR product for VRC26UCA HC was longer than that for the driver HC; the identities of the PCR products were confirmed with sequencing. In correlation with input DNA, the DC-PCR reaction yielded equal amounts of products for driver HC and VRC26UCA HC.

**Fig.S9.** PCR analysis of hybridomas derived from splenic B cells of the conditional model of VRC26UCA.

Panel **A** and **B** show RT-PCR amplification of driver HC **(A)** and VRC26UCA HC **(B)** transcripts from 8 hybridomas. The diagram above the gel image indicated the position of the PCR primers. Since these hybridomas secreted IgG1, the reverse primer was complementary to Cγ1. Panel **C** and **D** show PCR amplification of driver HC **(C)** and VRC26UCA HC **(D)** DNA from the same 8 hybridomas as shown in panels **A** and **B**. The diagram above the gel image indicated the position of the PCR primers. This experiment shows that the driver HC prevented the expression of VRC26UCA HC, and deletion of the driver variable region led to expression of VRC26UCA HC. The result validated the design of the conditional expression cassette.

**Table S1.** Summary of V_H_ replacement products isolated from a constitutive expression model of VRC26UCA.

This table listed V_H_ replacement products that were isolated from the hybridomas. The recombination took place between mouse V_H_ and D segments with the KI VRC26UCA HC on IgH^a^ allele from 129Sv mouse strain. The IgH locus from the 129Sv mouse strain has not been completely annotated. The V_H_ and D segments in the table were the best matches from the IMGT data bases. Since the analysis was intended to illustrate the structure of V_H_ replacement product, the precise identities of the gene segments were not critical.

## References

1. Burton DR & Hangartner L (2016) Broadly Neutralizing Antibodies to HIV and Their Role in Vaccine Design. Annual review of immunology 34:635–659.

2. Haynes BF & Mascola JR (2017) The quest for an antibody-based HIV vaccine. Immunological reviews 275(1):5–10.

3. Verkoczy L, Alt FW, & Tian M (2017) Human Ig knockin mice to study the development and regulation of HIV-1 broadly neutralizing antibodies. Immunological reviews 275(1):89–107.

4. Jung D, Giallourakis C, Mostoslavsky R, & Alt FW (2006) Mechanism and control of V(D)J recombination at the immunoglobulin heavy chain locus. Annual review of immunology 24:541–570.

5. Verkoczy L, et al. (2011) Rescue of HIV-1 broad neutralizing antibody-expressing B cells in 2F5 VH x VL knockin mice reveals multiple tolerance controls. Journal of immunology (Baltimore, Md. : 1950) 187(7):3785–3797.

6. Verkoczy L, et al. (2010) Autoreactivity in an HIV-1 broadly reactive neutralizing antibody variable region heavy chain induces immunologic tolerance. Proceedings of the National Academy of Sciences of the United States of America 107(1):181–186.

7. Chen Y, et al. (2013) Common tolerance mechanisms, but distinct cross-reactivities associated with gp41 and lipids, limit production of HIV-1 broad neutralizing antibodies 2F5 and 4E10. Journal of immunology (Baltimore, Md. : 1950) 191(3):1260–1275.

8. Doyle-Cooper C, et al. (2013) Immune tolerance negatively regulates B cells in knock-in mice expressing broadly neutralizing HIV antibody 4E10. Journal of immunology (Baltimore, Md. : 1950) 191(6):3186–3191.

9. McGuire AT, et al. (2016) Specifically modified Env immunogens activate B-cell precursors of broadly neutralizing HIV-1 antibodies in transgenic mice. Nature communications 7:10618.

10. Lin YC, et al. (2018) One-step CRISPR/Cas9 method for the rapid generation of human antibody heavy chain knock-in mice. The EMBO journal 37(18).

11. Finney J & Kelsoe G (2018) Poly- and autoreactivity of HIV-1 bNAbs: implications for vaccine design. Retrovirology 15(1):53.

12. Bonsignori M, et al. (2014) An autoreactive antibody from an SLE/HIV-1 individual broadly neutralizes HIV-1. The Journal of clinical investigation 124(4):1835–1843.

13. Haynes BF, et al. (2005) Cardiolipin polyspecific autoreactivity in two broadly neutralizing HIV-1 antibodies. Science (New York, N.Y.) 308(5730):1906–1908.

14. Liu M, et al. (2015) Polyreactivity and autoreactivity among HIV-1 antibodies. Journal of virology 89(1):784–798.

15. Yang G, et al. (2013) Identification of autoantigens recognized by the 2F5 and 4E10 broadly neutralizing HIV-1 antibodies. The Journal of experimental medicine 210(2):241–256.

16. Finney J, et al. (2019) Cross-Reactivity to Kynureninase Tolerizes B Cells That Express the HIV-1 Broadly Neutralizing Antibody 2F5. Journal of immunology (Baltimore, Md. : 1950).

17. Chen C, Nagy Z, Prak EL, & Weigert M (1995) Immunoglobulin heavy chain gene replacement: a mechanism of receptor editing. Immunity 3(6):747–755.

18. Chen C, et al. (1994) Deletion and editing of B cells that express antibodies to DNA. Journal of immunology (Baltimore, Md. : 1950) 152(4):1970–1982.

19. Tiegs SL, Russell DM, & Nemazee D (1993) Receptor editing in self-reactive bone marrow B cells. The Journal of experimental medicine 177(4):1009–1020.

20. Gay D, Saunders T, Camper S, & Weigert M (1993) Receptor editing: an approach by autoreactive B cells to escape tolerance. The Journal of experimental medicine 177(4):999–1008.

21. Prak EL & Weigert M (1995) Light chain replacement: a new model for antibody gene rearrangement. The Journal of experimental medicine 182(2):541–548.

22. Radic MZ, Erikson J, Litwin S, & Weigert M (1993) B lymphocytes may escape tolerance by revising their antigen receptors. The Journal of experimental medicine 177(4):1165–1173.

23. Adams E, Basten A, & Goodnow CC (1990) Intrinsic B-cell hyporesponsiveness accounts for self-tolerance in lysozyme/anti-lysozyme double-transgenic mice. Proceedings of the National Academy of Sciences of the United States of America 87(15):5687–5691.

24. Goodnow CC, et al. (1988) Altered immunoglobulin expression and functional silencing of self-reactive B lymphocytes in transgenic mice. Nature 334(6184):676–682.

25. Goodnow CC, Crosbie J, Jorgensen H, Brink RA, & Basten A (1989) Induction of self-tolerance in mature peripheral B lymphocytes. Nature 342(6248):385–391.

26. Hartley SB, et al. (1991) Elimination from peripheral lymphoid tissues of self-reactive B lymphocytes recognizing membrane-bound antigens. Nature 353(6346):765–769.

27. Nemazee DA & Burki K (1989) Clonal deletion of B lymphocytes in a transgenic mouse bearing anti-MHC class I antibody genes. Nature 337(6207):562–566.

28. Cyster JG & Goodnow CC (1995) Antigen-induced exclusion from follicles and anergy are separate and complementary processes that influence peripheral B cell fate. Immunity 3(6):691–701.

29. Cyster JG, Hartley SB, & Goodnow CC (1994) Competition for follicular niches excludes self-reactive cells from the recirculating B-cell repertoire. Nature 371(6496):389–395.

30. Kelsoe G & Haynes BF (2017) Host controls of HIV broadly neutralizing antibody development. Immunological reviews 275(1):79–88.

31. Doria-Rose NA, et al. (2016) New Member of the V1V2-Directed CAP256-VRC26 Lineage That Shows Increased Breadth and Exceptional Potency. Journal of virology 90(1):76–91.

32. Doria-Rose NA, et al. (2014) Developmental pathway for potent V1V2-directed HIV-neutralizing antibodies. Nature 509(7498):55–62.

33. Gorman J, et al. (2016) Structures of HIV-1 Env V1V2 with broadly neutralizing antibodies reveal commonalities that enable vaccine design. Nature structural & molecular biology 23(1):81–90.

34. DeKosky BJ, et al. (2015) In-depth determination and analysis of the human paired heavy- and light-chain antibody repertoire. Nature medicine 21(1):86–91.

35. Zemlin M, et al. (2003) Expressed murine and human CDR-H3 intervals of equal length exhibit distinct repertoires that differ in their amino acid composition and predicted range of structures. Journal of molecular biology 334(4):733–749.

36. Meffre E, et al. (2001) Immunoglobulin heavy chain expression shapes the B cell receptor repertoire in human B cell development. The Journal of clinical investigation 108(6):879–886.

37. Maruyama M, Lam KP, & Rajewsky K (2000) Memory B-cell persistence is independent of persisting immunizing antigen. Nature 407(6804):636–642.

38. Kavaler J, Caton AJ, Staudt LM, Schwartz D, & Gerhard W (1990) A set of closely related antibodies dominates the primary antibody response to the antigenic site CB of the A/PR/8/34 influenza virus hemagglutinin. Journal of immunology (Baltimore, Md. : 1950) 145(7):2312–2321.

39. Kraus M, Alimzhanov MB, Rajewsky N, & Rajewsky K (2004) Survival of resting mature B lymphocytes depends on BCR signaling via the Igalpha/beta heterodimer. Cell 117(6):787–800.

40. Pelanda R, Schaal S, Torres RM, & Rajewsky K (1996) A prematurely expressed Ig(kappa) transgene, but not V(kappa)J(kappa) gene segment targeted into the Ig(kappa) locus, can rescue B cell development in lambda5-deficient mice. Immunity 5(3):229–239.

41. Sonoda E, et al. (1997) B cell development under the condition of allelic inclusion. Immunity 6(3):225–233.

42. Bothwell AL, et al. (1981) Heavy chain variable region contribution to the NPb family of antibodies: somatic mutation evident in a gamma 2a variable region. Cell 24(3):625–637.

43. Reth M, Hammerling GJ, & Rajewsky K (1978) Analysis of the repertoire of anti-NP antibodies in C57BL/6 mice by cell fusion. I. Characterization of antibody families in the primary and hyperimmune response. European journal of immunology 8(6):393–400.

44. Ozato K, Mayer N, & Sachs DH (1980) Hybridoma cell lines secreting monoclonal antibodies to mouse H-2 and Ia antigens. Journal of immunology (Baltimore, Md. : 1950) 124(2):533–540.

45. Casellas R, et al. (1998) Ku80 is required for immunoglobulin isotype switching. The EMBO journal 17(8):2404–2411.

46. Taki S, Schwenk F, & Rajewsky K (1995) Rearrangement of upstream DH and VH genes to a rearranged immunoglobulin variable region gene inserted into the DQ52-JH region of the immunoglobulin heavy chain locus. European journal of immunology 25(7):1888–1896.

47. Hardy RR & Shinton SA (2004) Characterization of B lymphopoiesis in mouse bone marrow and spleen. Methods in molecular biology (Clifton, N.J.) 271:1–24.

48. Melchers F (2005) The pre-B-cell receptor: selector of fitting immunoglobulin heavy chains for the B-cell repertoire. Nature reviews. Immunology 5(7):578–584.

49. Martensson IL, et al. (2002) The pre-B cell receptor and its role in proliferation and Ig heavy chain allelic exclusion. Seminars in immunology 14(5):335–342.

50. Grawunder U, et al. (1995) Down-regulation of RAG1 and RAG2 gene expression in preB cells after functional immunoglobulin heavy chain rearrangement. Immunity 3(5):601–608.

51. Kitamura D, et al. (1992) A critical role of lambda 5 protein in B cell development. Cell 69(5):823–831.

52. Loffert D, Ehlich A, Muller W, & Rajewsky K (1996) Surrogate light chain expression is required to establish immunoglobulin heavy chain allelic exclusion during early B cell development. Immunity 4(2):133–144.

53. Papavasiliou F, Jankovic M, & Nussenzweig MC (1996) Surrogate or conventional light chains are required for membrane immunoglobulin mu to activate the precursor B cell transition. The Journal of experimental medicine 184(5):2025–2030.

54. Rolink A, et al. (1993) B cell development in mice with a defective lambda 5 gene. European journal of immunology 23(6):1284–1288.

55. Shimizu T, Mundt C, Licence S, Melchers F, & Martensson IL (2002) VpreB1/VpreB2/lambda 5 triple-deficient mice show impaired B cell development but functional allelic exclusion of the IgH locus. Journal of immunology (Baltimore, Md. : 1950) 168(12):6286–6293.

56. Kudo A & Melchers F (1987) A second gene, VpreB in the lambda 5 locus of the mouse, which appears to be selectively expressed in pre-B lymphocytes. The EMBO journal 6(8):2267–2272.

57. Sakaguchi N & Melchers F (1986) Lambda 5, a new light-chain-related locus selectively expressed in pre-B lymphocytes. Nature 324(6097):579–582.

58. Lam KP, Kuhn R, & Rajewsky K (1997) In vivo ablation of surface immunoglobulin on mature B cells by inducible gene targeting results in rapid cell death. Cell 90(6):1073–1083.

59. Li YS, Wasserman R, Hayakawa K, & Hardy RR (1996) Identification of the earliest B lineage stage in mouse bone marrow. Immunity 5(6):527–535.

60. Tian M, et al. (2016) Induction of HIV Neutralizing Antibody Lineages in Mice with Diverse Precursor Repertoires. Cell 166(6):1471–1484.e1418.

61. Bankovich AJ, et al. (2007) Structural insight into pre-B cell receptor function. Science (New York, N.Y.) 316(5822):291–294.

62. Rettig TA, Ward C, Bye BA, Pecaut MJ, & Chapes SK (2018) Characterization of the naive murine antibody repertoire using unamplified high-throughput sequencing. PloS one 13(1):e0190982.

63. Kelsoe G, Verkoczy L, & Haynes BF (2014) Immune System Regulation in the Induction of Broadly Neutralizing HIV-1 Antibodies. Vaccines 2(1):1–14.

64. Chen J, Lansford R, Stewart V, Young F, & Alt FW (1993) RAG-2-deficient blastocyst complementation: an assay of gene function in lymphocyte development. Proceedings of the National Academy of Sciences of the United States of America 90(10):4528–4532.

65. Shinkura R, et al. (2003) The influence of transcriptional orientation on endogenous switch region function. Nature immunology 4(5):435–441.

66. Harlow E, & Lane, D. (1988) Antibodies: a laboratory manual (Cold Spring Harbor Laboratory,Cold Spring Harbor, NY).

67. Tom R, Bisson L, & Durocher Y (2008) Transfection of HEK293-EBNA1 Cells in Suspension with Linear PEI for Production of Recombinant Proteins. CSH protocols 2008:pdb.prot4977.

