## Supplementary material for "Conditional Antibody Expression to Avoid Central B Cell Deletion in a Humanized HIV-1 Vaccine Mouse Models": Fig.S1-S9, Table S1

**Figure S1**

**A.**

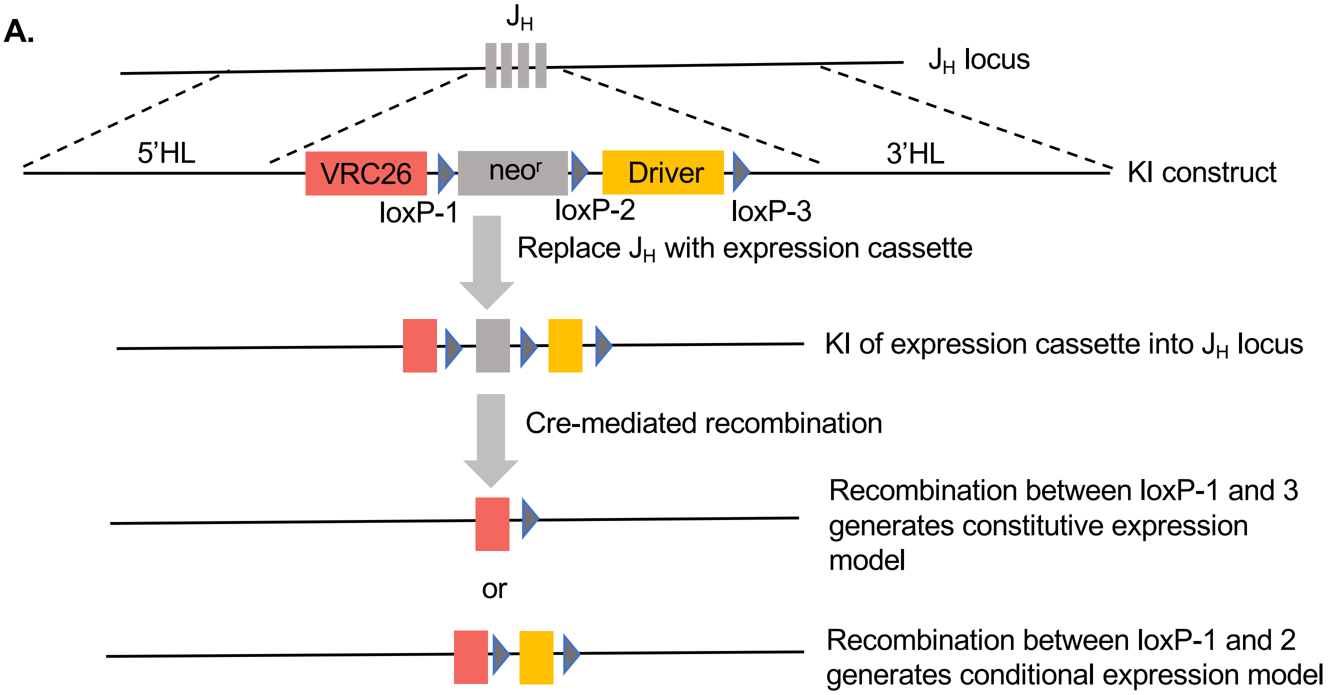


**B.**

| Digest | Probe | Germline | Constitutive model | Conditional model |
|---|---|---|---|---|
| EcoRV | $J_H$ 3' | 16,530bp | 10,808bp | 10,968bp |
| NheI | $J_H$ 5' | 5,509bp | 13,863bp | 7,729bp |
| EcoRV+SpeI | Jκ 3' | 8,653bp | 5,502bp | 5,502bp |
| PstI | Jκ 5' | 7,006bp | 5,263bp | 5,263bp |

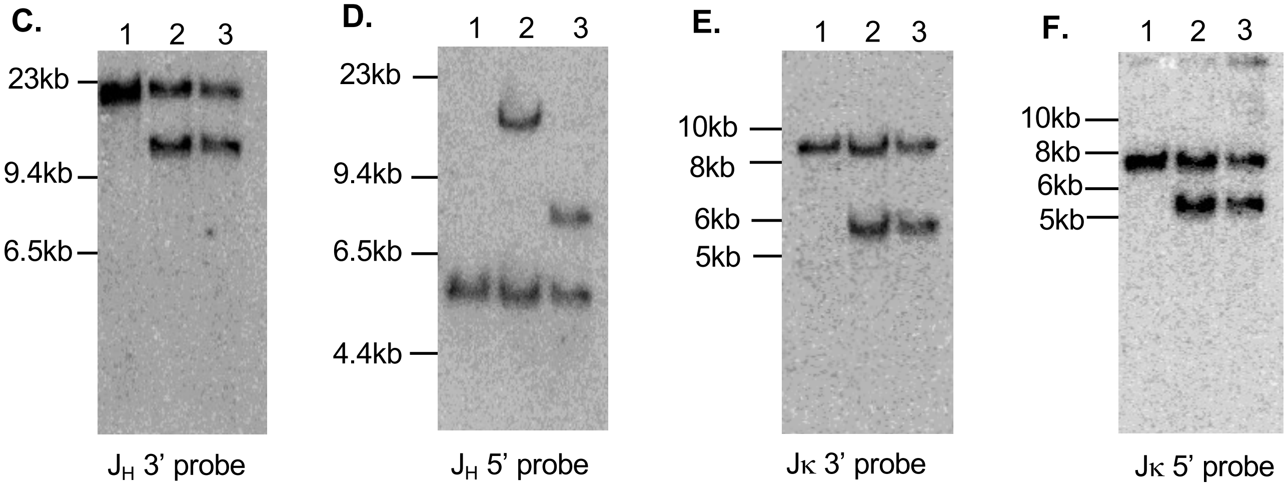

### Figure S2

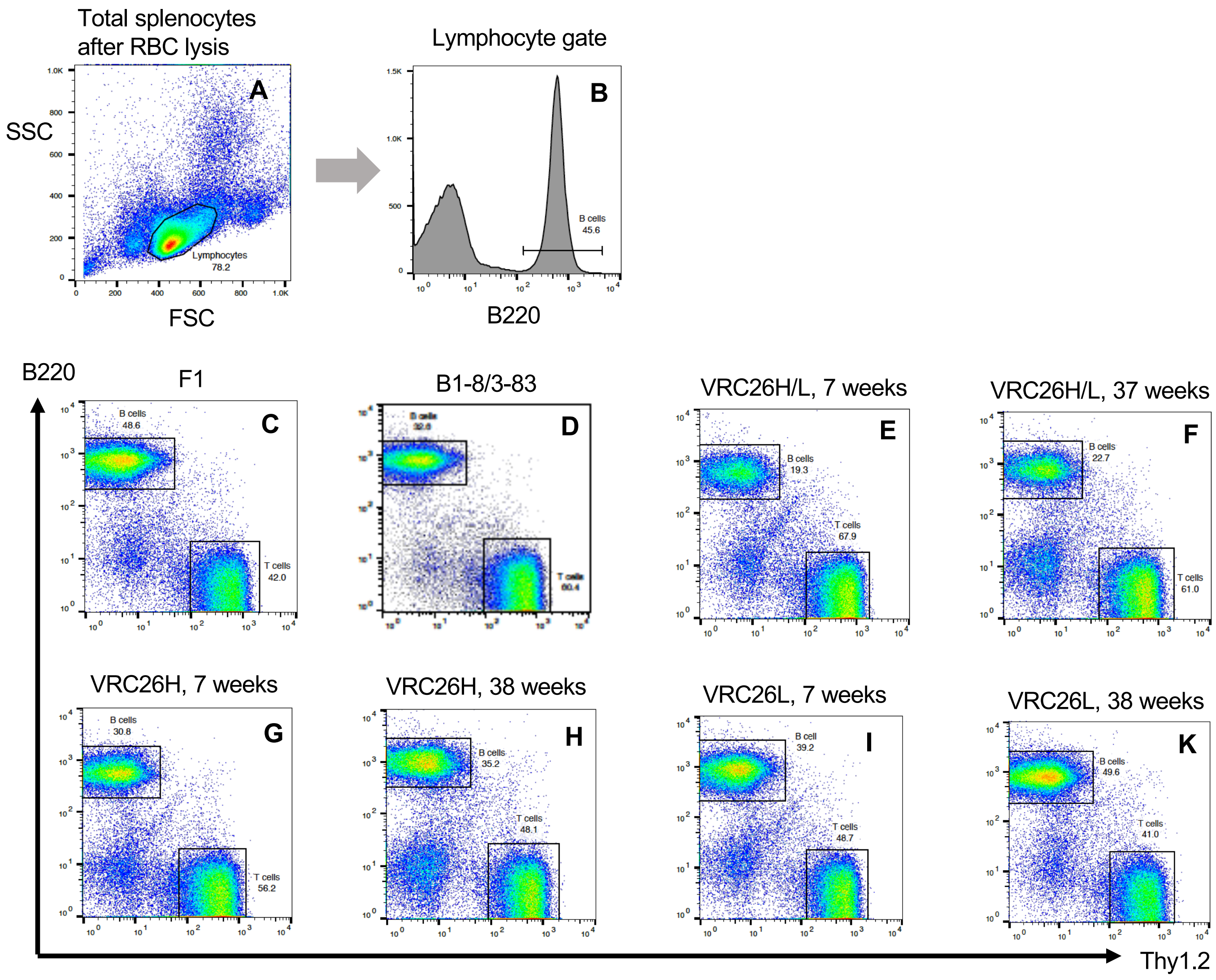

Total splenocytes after RBC lysis
Lymphocyte gate
A
B
SSC
FSC
Lymphocytes 78.2
B220
B cells 45.6
B220
Thy1.2
F1
B1-8/3-83
VRC26H/L, 7 weeks
VRC26H/L, 37 weeks
VRC26H, 7 weeks
VRC26H, 38 weeks
VRC26L, 7 weeks
VRC26L, 38 weeks
C
B cells 48.6
T cells 42.0
D
E
B cells 19.3
T cells 67.9
F
B cells 22.7
T cells 61.0
G
B cells 30.8
T cells 56.2
H
B cells 35.2
T cells 48.1
I
B cell 39.2
T cells 48.7
K
B cells 49.6
T cells 41.0

**Figure S3**

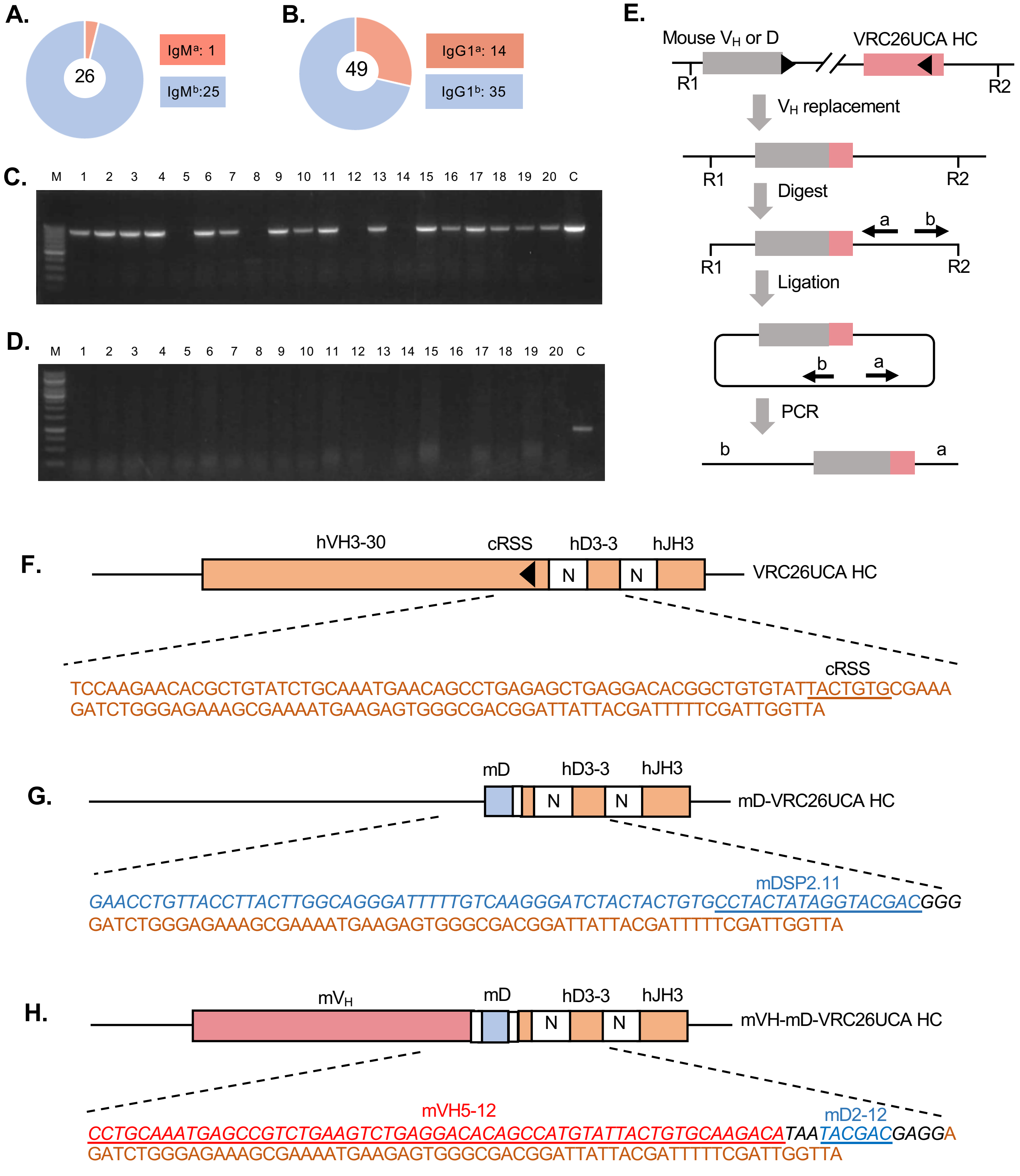

A.
26
IgMa: 1
IgMb:25
B.
49
IgG1a: 14
IgG1b: 35
C.
M 1 2 3 4 5 6 7 8 9 10 11 12 13 14 15 16 17 18 19 20 C
D.
M 1 2 3 4 5 6 7 8 9 10 11 12 13 14 15 16 17 18 19 20 C
E.
Mouse VH or D
VRC26UCA HC
R1
R2
VH replacement
Digest
a
b
Ligation
PCR
F.
hVH3-30
cRSS
hD3-3
hJH3
N
VRC26UCA HC
TCCAAGAACACGCTGTATCTGCAAATGAACAGCCTGAGAGCTGAGGACACGGCTGTGTATTACTGTGCGAAA
GATCTGGGAGAAAGCGAAAATGAAGAGTGGGCGACGGATTATTACGATTTTTCGATTGGTTA
G.
mD
mD-VRC26UCA HC
mDSP2.11
GAACCTGTTACCTTACTTGGCAGGGATTTTTGTCAAGGGATCTACTACTGTGCCTACTATAGGTACGACGGG
GATCTGGGAGAAAGCGAAAATGAAGAGTGGGCGACGGATTATTACGATTTTTCGATTGGTTA
H.
mVH
mVH-mD-VRC26UCA HC
mVH5-12
mD2-12
CCTGCAAATGAGCCGTCTGAAGTCTGAGGACACAGCCATGTATTACTGTGCAAGACATAATACGACGAGGA
GATCTGGGAGAAAGCGAAAATGAAGAGTGGGCGACGGATTATTACGATTTTTCGATTGGTTA

#### Figure S4

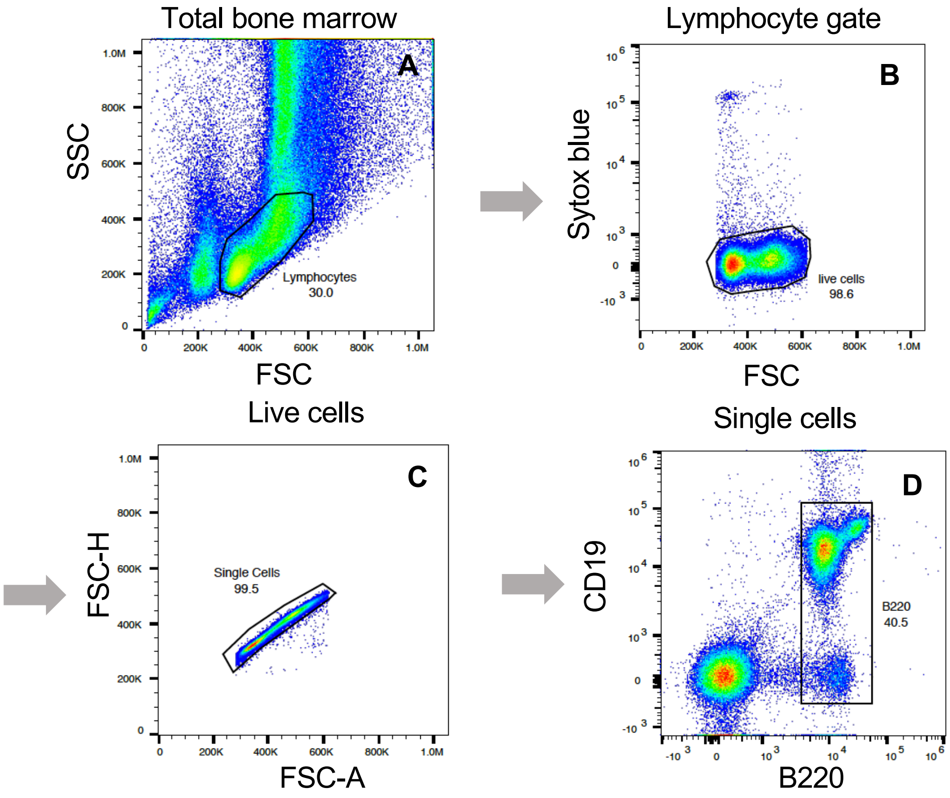
Total bone marrow
A
SSC
FSC
Lymphocytes
30.0
Lymphocyte gate
B
Sytox blue
FSC
live cells
98.6
Live cells
C
FSC-H
FSC-A
Single Cells
99.5
Single cells
D
CD19
B220
B220
40.5

#### Figure S5

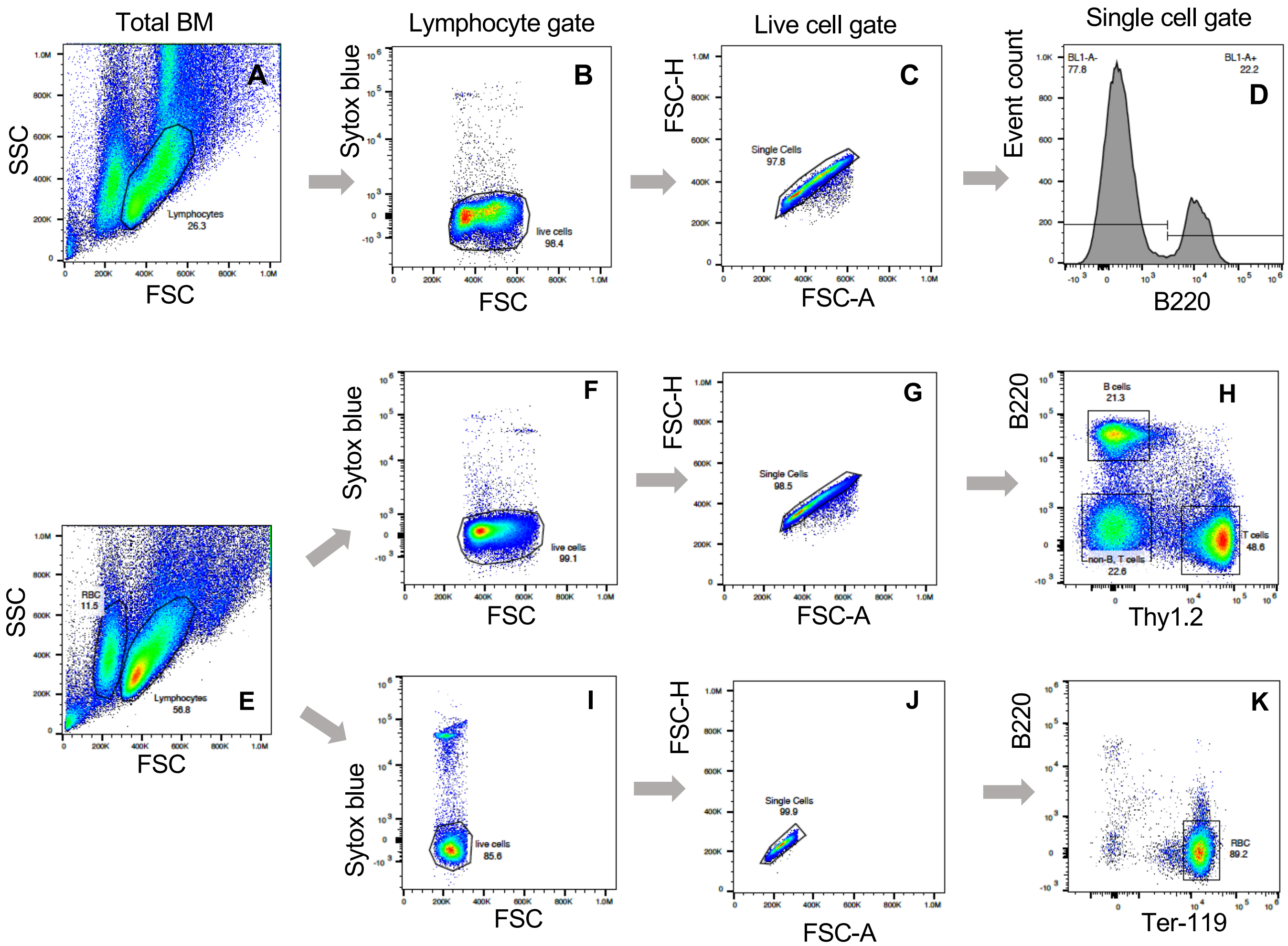

Total BM
Lymphocyte gate
Live cell gate
Single cell gate
A
B
C
D
E
F
G
H
I
J
K
SSC
FSC
Lymphocytes
26.3
Sytox blue
live cells
98.4
FSC-H
FSC-A
Single Cells
97.8
Event count
BL1-A-
77.8
BL1-A+
22.2
B220
RBC
11.5
Lymphocytes
56.8
live cells
99.1
Single Cells
98.5
B cells
21.3
T cells
48.6
non-B, T cells
22.6
Thy1.2
live cells
85.6
Single Cells
99.9
RBC
89.2
Ter-119

#### Figure S6

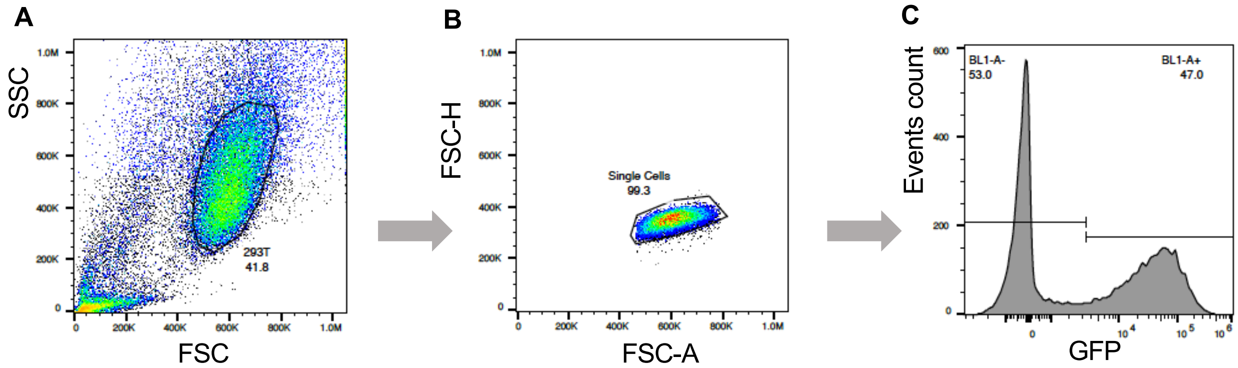


**D. GFP+ cells (%) in HC/mVpreB/mλ5 transfection**

| | Driver | VRC26 | VHB1-8 |
|---|---|---|---|
| Transfection 1 | 47.0 | 39.6 | 44.0 |
| Transfection 2 | 47.3 | 38.2 | 43.1 |
| Transfection 3 | 44.0 | 43.1 | 41.7 |
| Average | 46.1 | 40.3 | 42.9 |

**E. GFP+ cells (%) in HC/LC transfection**

| | Driver | VRC26 | VHB1-8 |
|---|---|---|---|
| Transfection 1 | 54.4 | 49.1 | 48.9 |
| Transfection 2 | 47.7 | 50.7 | 51.4 |
| Transfection 3 | 47.7 | 48.0 | 48.8 |
| Average | 49.9 | 49.3 | 49.7 |

**F. GFP+ cells (%) in HC/hVpreB/hλ5 transfection**

| | Driver | VRC26 |
|---|---|---|
| Transfection 1 | 52.2 | 52.1 |
| Transfection 2 | 49.0 | 49.9 |
| Transfection 3 | 54.2 | 52.7 |
| Average | 51.8 | 51.6 |

**G. GFP+ cells (%) in HC/hVpreB/mλ5 transfection**

| | Driver | VRC26 |
|---|---|---|
| Transfection 1 | 40.9 | 38.0 |
| Transfection 2 | 40.7 | 40.5 |
| Transfection 3 | 40.1 | 38.0 |
| Average | 40.6 | 38.8 |

**H. GFP+ cells (%) in HC/mVpreB/hλ5 transfection**

| | Driver | VRC26 |
|---|---|---|
| Transfection 1 | 53.1 | 55.0 |
| Transfection 2 | 54.8 | 52.0 |
| Transfection 3 | 53.7 | 52.0 |
| Average | 53.9 | 53.0 |

**Figures S7**

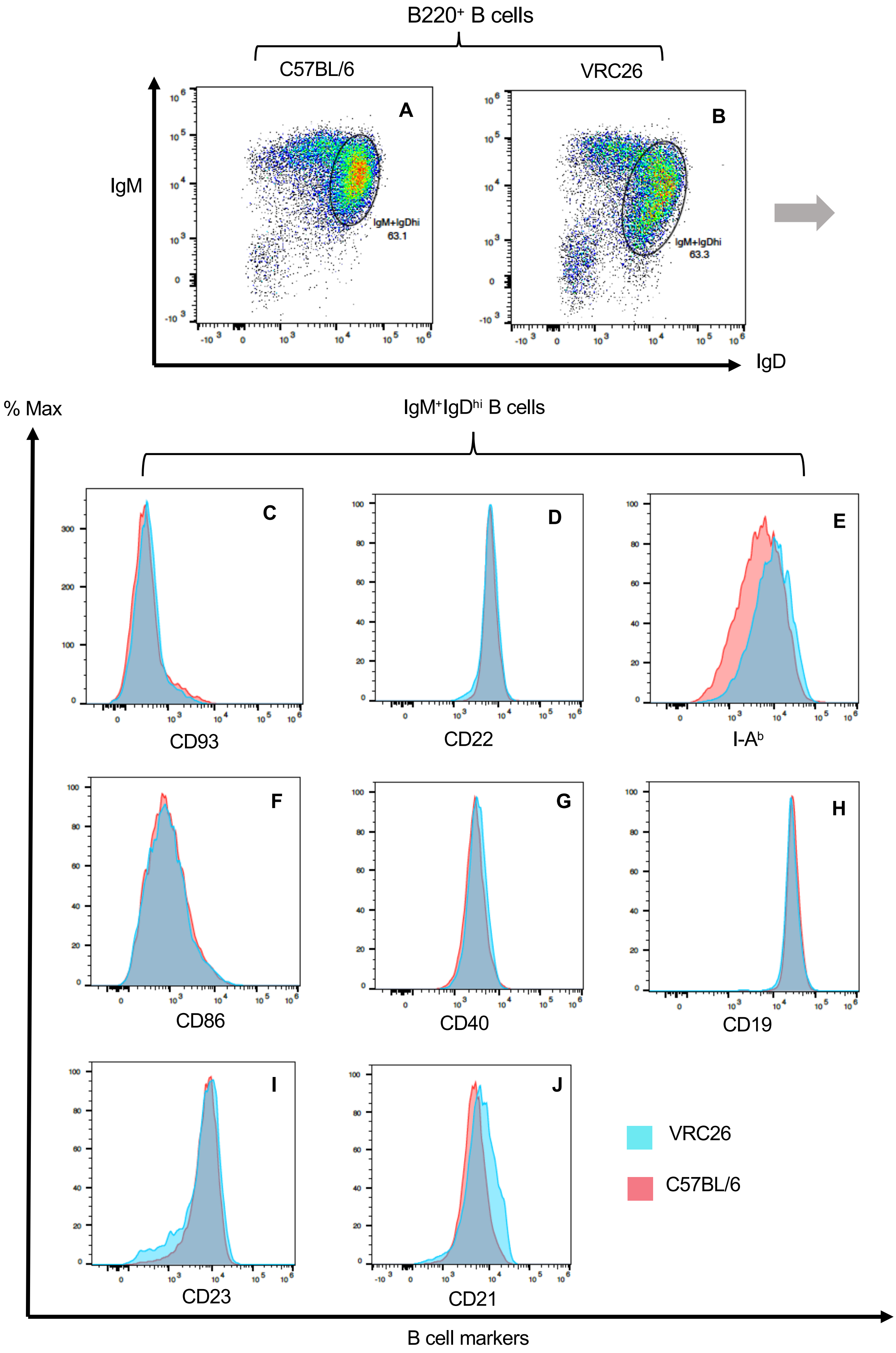
B220⁺ B cells
C57BL/6
VRC26
A
B
IgM
IgD
IgM+IgDhi
63.1
IgM+IgDhi
63.3
% Max
IgM⁺IgDʰⁱ B cells
C
D
E
CD93
CD22
I-Aᵇ
F
G
H
CD86
CD40
CD19
I
J
CD23
CD21
VRC26
C57BL/6
B cell markers

**Figure S8**

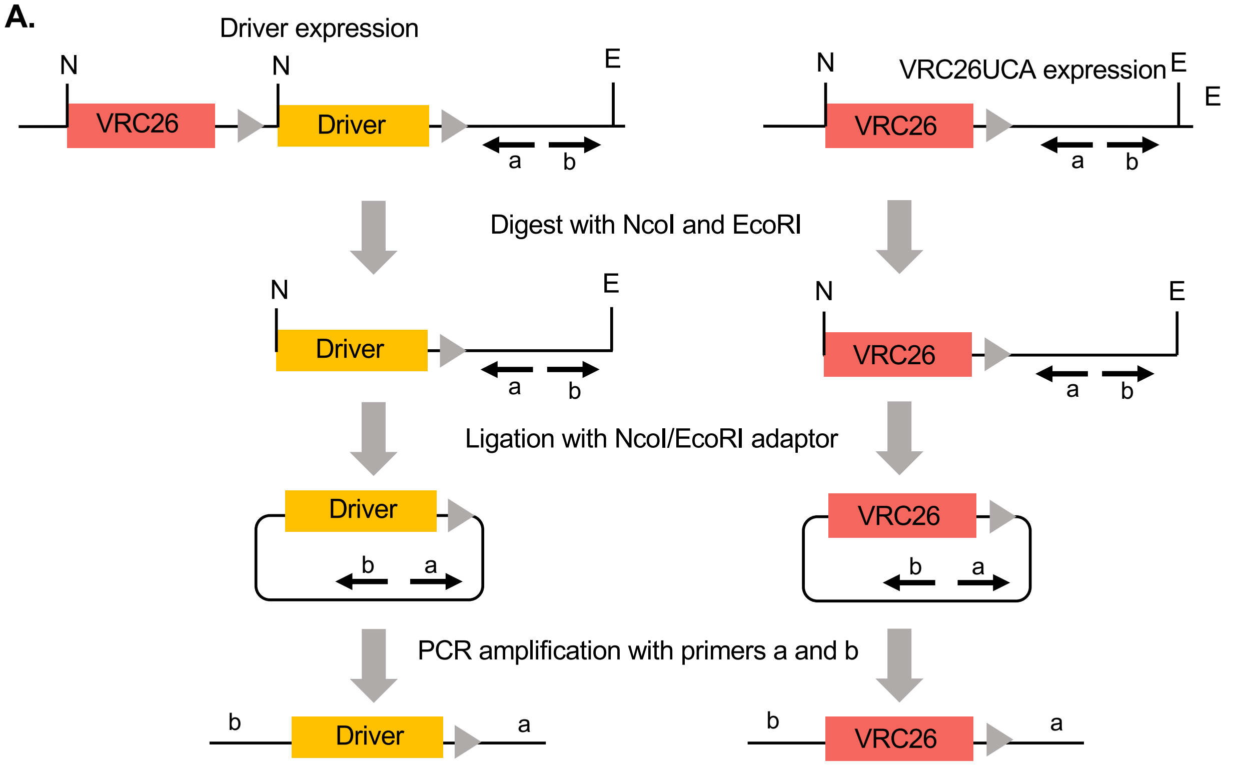
A.
Driver expression
N
N
E
VRC26
Driver
a
b
N
VRC26UCA expression
E
E
VRC26
a
b
Digest with NcoI and EcoRI
N
E
Driver
a
b
N
E
VRC26
a
b
Ligation with NcoI/EcoRI adaptor
Driver
b
a
VRC26
b
a
PCR amplification with primers a and b
b
Driver
a
b
VRC26
a

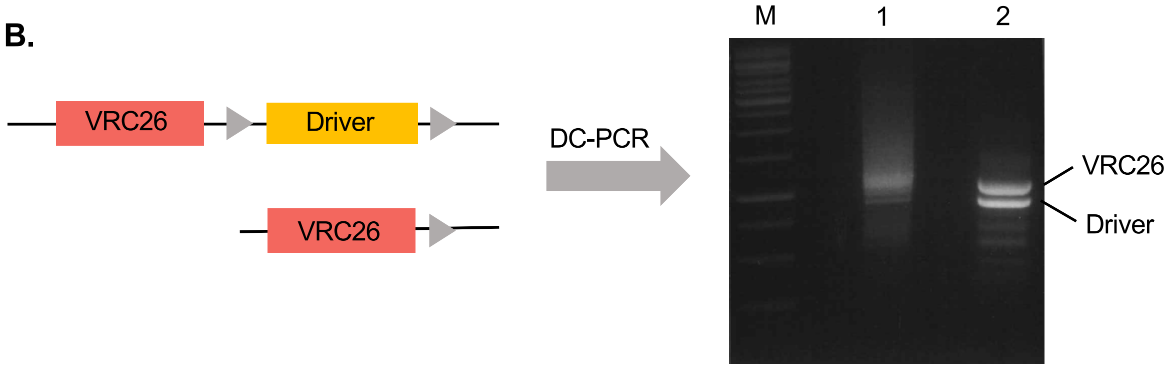
B.
VRC26
Driver
VRC26
DC-PCR
M
1
2
VRC26
Driver

**Figure S9**

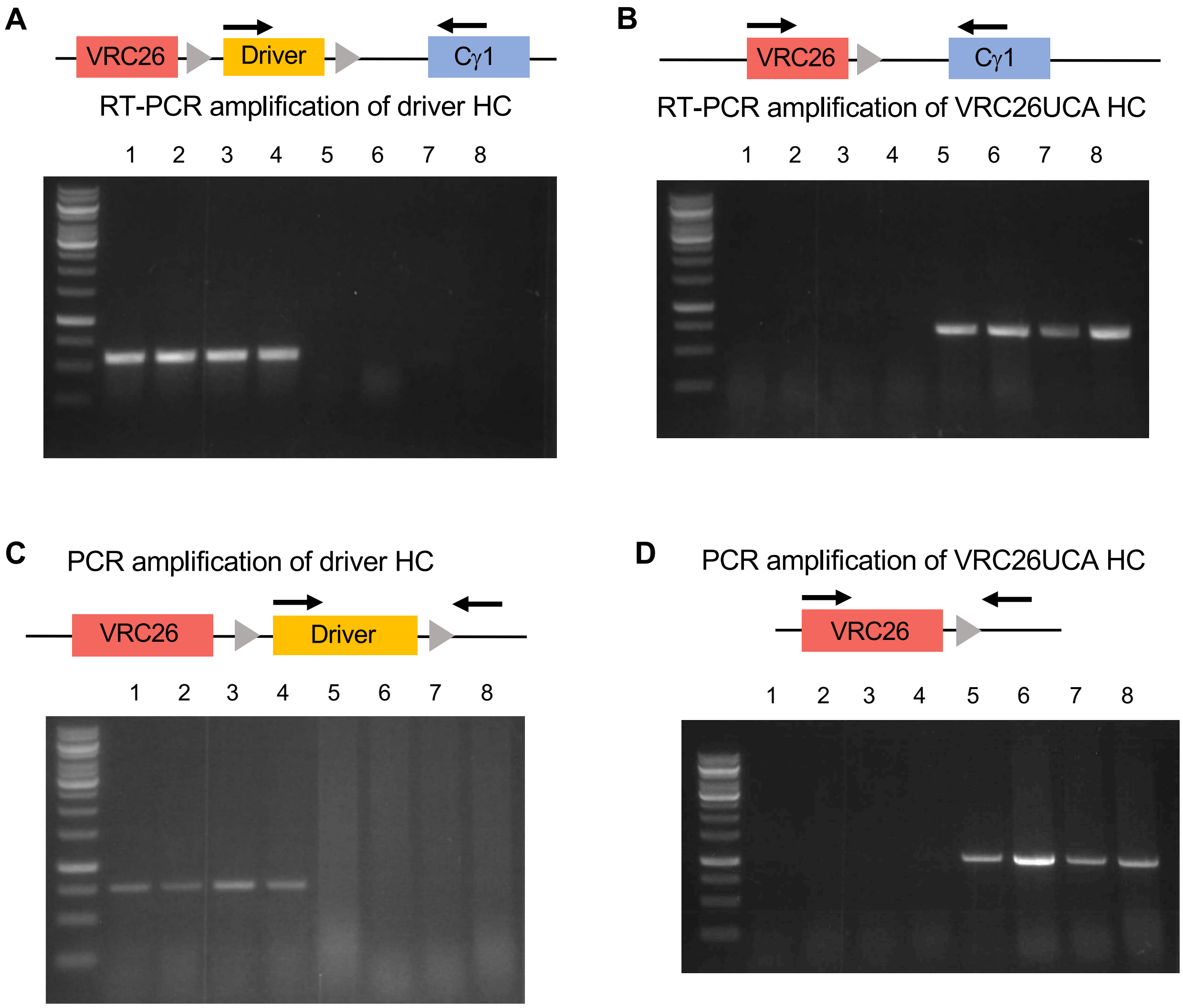

A
VRC26
Driver
Cγ1
RT-PCR amplification of driver HC
1 2 3 4 5 6 7 8
B
VRC26
Cγ1
RT-PCR amplification of VRC26UCA HC
1 2 3 4 5 6 7 8
C
PCR amplification of driver HC
VRC26
Driver
1 2 3 4 5 6 7 8
D
PCR amplification of VRC26UCA HC
VRC26
1 2 3 4 5 6 7 8

**Table S1**

Summary of VH replacement events in hybridomas from VRC26UCA KI mouse

| Hybridoma clone | Mouse VH | Mouse D | P/NP * | Antibody |
|---|---|---|---|---|
| 1 | - | IGHD2-10*02 | NP, no VH | IgMb |
| 2 | - | IGHD2-3*01 | NP, no VH | IgMb |
| 3 | - | IGHD2-14*01 | NP, no VH | IgMb |
| 4 | - | IGHD2-10*02 | NP, no VH | IgMb |
| 5 | - | IGHD2-10*02 | NP, no VH | IgMb |
| 6 | - | DSP2.8 | NP, no VH | IgG1b |
| 7 | VH5-12*02 | D2-12*01 | NP, out of frame | IgMb |
| 8 | IGHV5-6-5*01 | IGHD2-2*01 | NP, out of frame | IgMb |
| 9 | IGHV1S81*02 | IGHD2-1*01 | NP, out of frame | IgMb |
| 10 | IGHV2-7*01 | - | NP, out of frame | IgMb |
| 11 | IGHV8-2*01 | IGHD2-14*01 | NP, out of frame | IgG1b |
| 12 | IGHV9-2*01 | IGHD2-14*01 | NP, inframe with stop codon | IgMb |
| 13 | IGHV-69*02 | - | P | IgG1b |
| 14 | IGHV1-26*01# | IGHD2-3*01 | P | IgMa |
| 15 | IGHV8-12*01 | IGH2-4*01 | P | IgG1a |

* P, productive; NP, non-productive.
